# Diffusing protein binders to intrinsically disordered proteins

**DOI:** 10.1101/2024.07.16.603789

**Authors:** Caixuan Liu, Kejia Wu, Hojun Choi, Hannah Han, Xulie Zhang, Joseph L. Watson, Sara Shijo, Asim K. Bera, Alex Kang, Evans Brackenbrough, Brian Coventry, Derrick R. Hick, Andrew N. Hoofnagle, Ping Zhu, Xingting Li, Justin Decarreau, Stacey R. Gerben, Wei Yang, Xinru Wang, Mila Lamp, Analisa Murray, Magnus Bauer, David Baker

## Abstract

Proteins which bind intrinsically disordered proteins (IDPs) and intrinsically disordered regions (IDRs) with high affinity and specificity could have considerable utility for therapeutic and diagnostic applications. However, a general methodology for targeting IDPs/IDRs has yet to be developed. Here, we show that starting only from the target sequence of the input, and freely sampling both target and binding protein conformation, RFdiffusion can generate binders to IDPs and IDRs in a wide range of conformations. We use this approach to generate binders to the IDPs Amylin, C-peptide and VP48 in a range of conformations with Kds in the 3 -100nM range. The Amylin binder inhibits amyloid fibril formation and dissociates existing fibers, and enables enrichment of amylin for mass spectrometry-based detection. For the IDRs G3bp1, common gamma chain (IL2RG) and prion, we diffused binders to beta strand conformations of the targets, obtaining 10 to 100 nM affinity. The IL2RG binder colocalizes with the receptor in cells, enabling new approaches to modulating IL2 signaling. Our approach should be widely useful for creating binders to flexible IDPs/IDRs spanning a wide range of intrinsic conformational preferences.

## Main

IDPs and IDPRs (structured proteins with intrinsically disordered regions) are abundant in nature and carry out important biological functions without adopting a single well-defined structure, and hence are well established biomarkers in clinical care and biomedical research (Fig. 1a). Designing binders specific for disordered regions could be valuable for clinical diagnosis, therapeutic development, and scientific research^1–4^. Current methods largely rely on antibodies, which have limitations such as high production costs, reproducibility, and complex engineering requirements^5,6^; the dynamic nature of disordered proteins can also complicate the elicitation of antibodies^7,8^. Computational protein design has created binders of peptides in extended beta strand^9,10^, helical^11^, and polyproline II conformations^12^. While powerful, these methods require prespecification of the target peptide geometry, which can be limiting because the optimal conformation given both the intrinsic sequence biases of the peptide, and the opportunities for making high affinity interactions, may be quite irregular.

**Figure. 1.**
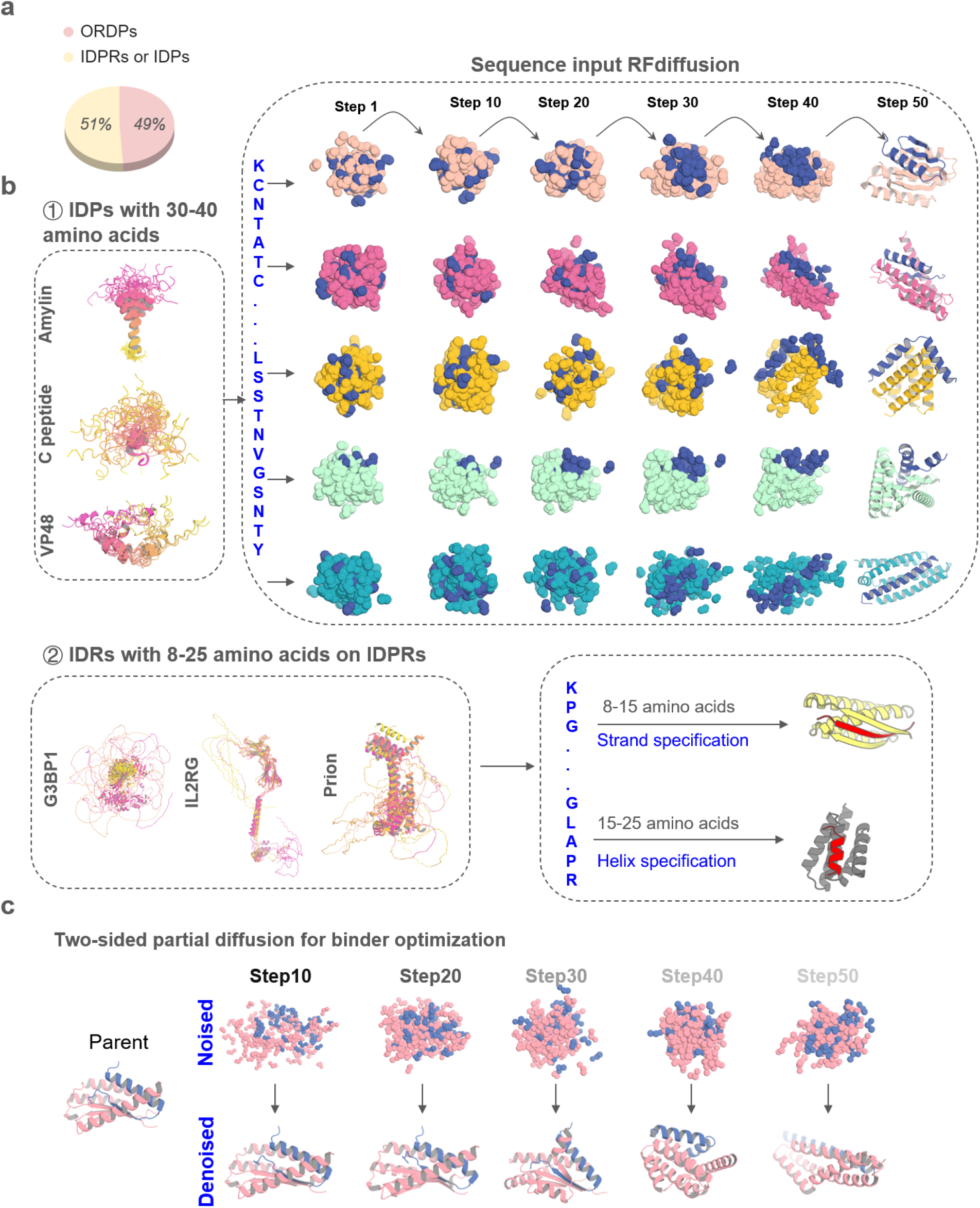
Design strategies for binding conformational flexible peptides. a, Frequency of ORDPs (ordered proteins), IDPRs /IDPs (intrinsically disordered proteins) in the human proteome^41^. **b,** ① Left, the NMR structure of Amylin (PDBID: 2KB8), C peptide (PDBID: 1T0C), the predicted structures of VP48 by five AlphaFold models^22^. The 5 predicted structures of VP48 are aligned together, revealing the flexibility of the intrinsically disordered protein. Right, Diffusion models for proteins are trained to recover noised protein structures and to generate new structures by reversing the corruption process through iterative denoising of initially random noise into a realistic structure. Here, A modified version of RFdiffusion was trained on two chain systems from the PDB to permit the design of protein binders to targets, for which only the sequence of the target was specified. The fine-tuned was found to generate binders to peptides in finely varying helix conformationsWith solely sequence input. ② Left, the predicted structures of G3BP1, IL2RG and prion by five AlphaFold models^22^. Right, A modified version of RFdiffusion was trained, allowing for specification of the secondary structure of a region, along with its sequence (See Method). When provided with the same target sequence input but different secondary structure specifications (helix or strand), the resulting conformations of the target could vary. **c,** Top: two sided partial diffusion. RFdiffusion is used to denoise a randomly noised starting parent design for both target and binder ; varying the extent by different noised step of initial noising (top row) enables control over the extent of introduced structural variation (bottom row; colours, new designs; grey, parent design).

We sought to develop a general approach to design high-affinity binders for intrinsically disordered proteins that starts from the target sequence alone and does not require prespecification of the target geometry (Fig. 1b ①). We reasoned that a version of RFdiffusion trained on two chain systems from the PDB, noising the structure on one and providing only the sequence on the second, could have such capability. This was used previously to generate binders to bioactive peptide hormones restricted to helical conformations^11^; here we begin by investigating the application of the approach to IDPs in a much broader range of conformations (the sequences of many targets are not compatible with uninterrupted helical conformations). To target shorter IDRs, we reasoned that strand pairing, as employed by Sahtoe et al using Rosetta^13^, coupled with RFdiffusion^14^ to sample the many different possible variations of strand conformation, could provide a general approach to maximizing interactions over a short region since backbone-backbone hydrogen bonds contribute to binding energy in addition to sidechain-sidechain interactions (Fig. 1b ②).

We first experimented with designing binders to the human islet amyloid polypeptide (hIAPP), also known as amylin, a 37-residue hormone co-secreted with insulin by pancreatic islet β-cells to modulate glucose levels^15,16^. Cysteine residues 2 and 7 form disulfide bridge which is critical for the full biological activity of amylin^15^. NMR studies conducted in lipid environments or under SDS micelle binding conditions have indicated helical propensity in Amylin fragments^17,18^; the overall structure appears to be intrinsically disordered^19,20^.

We employed the flexible target fine-tuned RFdiffusion to design binders against Amylin using only the Amylin sequence as input – the structure of the binding protein, the Amylin conformation, and the binding mode are entirely unspecified. Starting from the amino acid sequence of Amylin, RFdiffusion generated complexes encompassing a variety of conformations for both peptides and binders. Representative design trajectories are shown in Supplementary Video 1; starting from a random distribution of residues of both Amylin and binder; in sequential denoising steps, the Amylin adopts different conformations while the binder residue distribution shifts to surround Amylin and progressively organizes into a folded structure which cradles nearly the entire surface of the peptide (Fig. 1b①). The resulting library of backbones were sequence designed using ProteinMPNN^21^, and filtered using AlphaFold2 (AF2)^22^ for the monomer conformation and AF2 initial guess for the complex^23^.

We obtained synthetic genes encoding 96 designs binding amylin in a variety of conformations, expressed the proteins in *E.Coli*, and purified them using immobilized metal ion affinity chromatography (IMAC). Amylin binding affinities determined using bio-layer interferometry (BLI) ranged from 100 nM to 454 nM (Supplementary Fig. 1a). Since binders to peptides in entirely helical conformations have been studied^11^, here we focused on other geometries. To optimize the binding affinity of initial hits to αβ, αβL, and αα conformations, we implemented a two sided partial diffusion approach (see Methods; in contrast to one sided partial diffusion which only diversifies the binder conformation and keeps the target fixed, two sided partial diffusion allows simultaneous conformation changes of both target and binder which leads to broader sampling (Fig. 1c, Supplementary Fig. 2a)). We carried out 5,000 two sided diffusion trajectories from initial designs noised over 5 to 20 steps (complete randomization corresponds to 50 steps), and found that this yielded designs with generally better metrics than one sided diffusion likely because the peptide conformation can adapt to that of binder resulting in greater shape complementarity and more extensive interactions (Supplementary Fig. 2). We obtained synthetic genes encoding the 174 resulting designs with the best metrics that span amylin conformations in the αβ, αβL, and αα conformations. 107 out of 174 refined designs bound Amylin; the highest affinity binders (Amylin-68nαβ, Amylin-36αβ, Amylin-75αα and Amylin-22αβL) which bind Amylin in different conformations, have affinities of 3.8 nM, 10 nM, 15 nM and 100 nM, respectively (Fig. 2a-d). While the Amylin adopts very different conformations in different designs, the diffusion process was able to maintain the disulfide bond, key to amylin function, in all designs^15^ (Fig. 2a-d). Circular dichroism studies showed that all four binders were largely helical as designed and thermostable up to 95 °C (Supplementary Fig. 1b)

**Figure.2.**
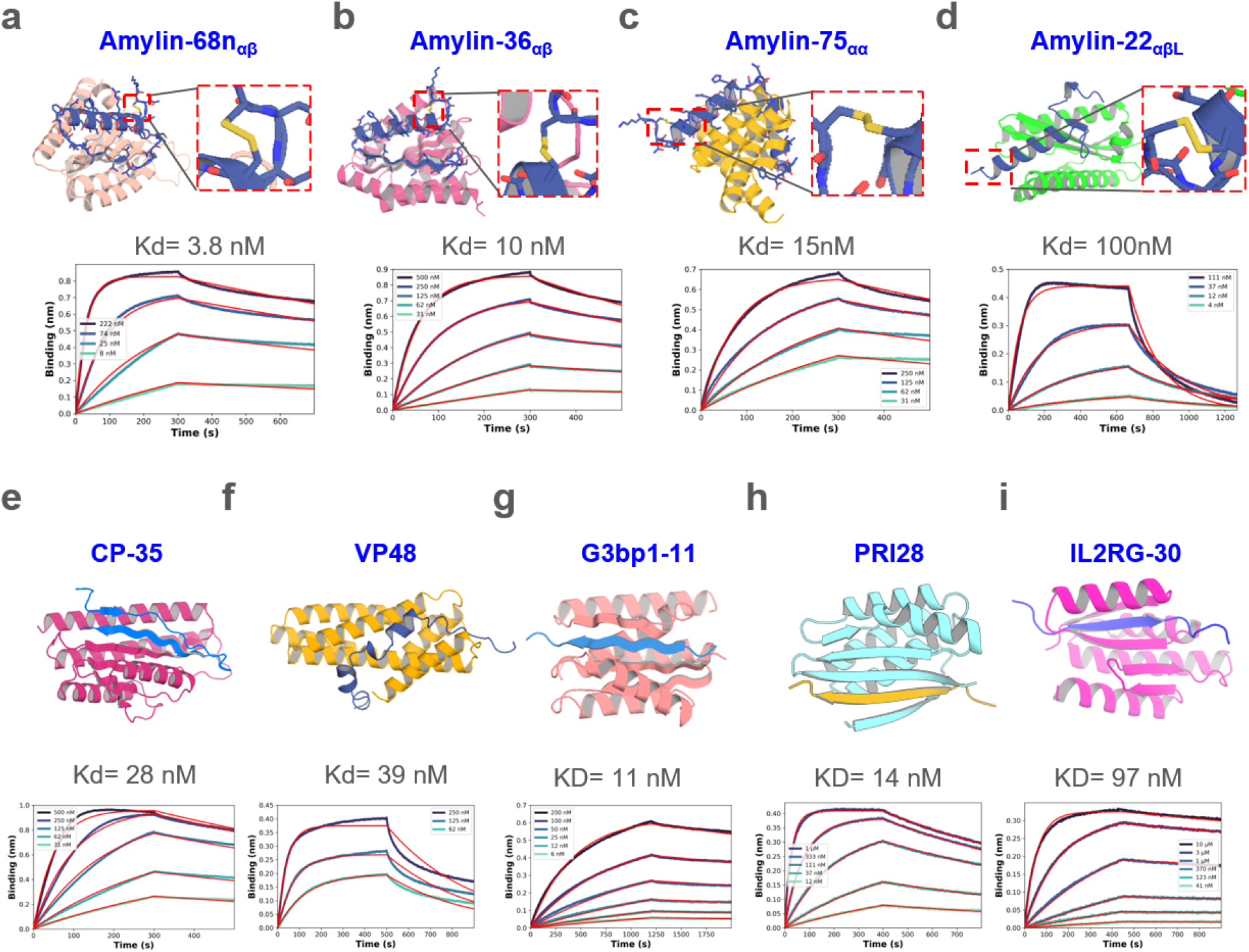
Design of disordered region binder. **a-d,** Binder design of Amylin using sequence input diffusion. Top, from left to right, design model of Amylin and its binder Amylin-68nαβ, Amylin-36αβ, Amylin-75αα and Amylin-22αβL, respectively. The secondary structure of Amylin is indicated in the subscript of the binder’s name. For each of the designs, the Amylin disulfide bonds between 2nd Cysteine and 7th Cysteine were retained well. Bottom, from left to right, the BLI measurement indicated that the binding affinity between Amylin-68nαβ, Amylin-36αβ, Amylin-75αα, Amylin-22αβL and Amylin are 3.8, 10, 15, 100 nM respectively. **e-f,** Binder design of CP and VP48 using sequence input diffusion, the binder affinity of CP and VP48 are 28 and 39 nM, respectively. **g-i,** Binder design using strand specification. Top, from left to right, design model of G3BP1^RBD^, prion and IL2RG and their binders G3bp1-11, PRI28 and IL2RG-30. Bottom, the BLI measurement indicated that the binding affinity of G3bp1-11, PRI28 and IL2RG-30 binders are 11, 14 and 97 nM, respectively.

C-peptide is a 31 residue peptide secreted by islet β cells that is made from the same precursor – proinsulin – as insulin^24^. Measurement of plasma C-peptide levels is important for accurate classification and diagnosis of type I and type II diabetes^25^. We carried out sequence-input diffusion with C-peptide allowed to sample diverse conformations (Supplementary Fig. 3a). Of 96 designs tested, one in which the C-peptide forms a long strand, followed by a long dynamic loop and a small strand paired with the long strand had weak binding affinity (Supplementary Fig. 3b-c). This design had more hydrogen bonds between target and binder (13) than all but 5 of the 96 designs (Supplementary Fig. 3d), and we hypothesized that this was important for binding. To optimize the initial hit to improve binding affinity, we again used two sided partial diffusion and included the number of hydrogen bonds in filtering. Screening with BLI revealed a much higher success rate, with six designs binding C peptide with better than 100nM binding affinity; the highest affinity binder (CP-35) had a Kd of 28nM (Fig. 2e). Circular dichroism studies showed that CP-35 was largely helical, consistent with the design model, and thermostable up to 95 °C (Supplementary Fig.3e).

We next chose to target VP48 (39 amino acid), a potent activator of transcription^26^. In a first round of 30,000 unconstrained RFdiffusion trajectories, the most enriched conformations after filtering contained substantial secondary structure as in the above cases. To explore binding to more loop-containing conformations, we filtered these designs based on target backbone conformation and a relatively loose PAE cutoff (PAE <16); within this pool, 20 designs were manually selected and further optimized by iterative partial diffusion and backbone extension (see Methods). Of 95 designs tested, 2 showed binding at 2 μM by BLI with the highest affinity 750nM for a design with the VP48 in a conformation with three short helical fragments connected with relatively long loops. Further partial diffusion optimization yielded a design with a Kd of 39nM (Fig. 2f), that again was thermostable up to 95C (Supplementary Fig. 3f).

## Targeting shorter IDRs using beta strand interactions

Consistent with the observations of Sahtoe et al using the non-deep learning Rosetta method^13^, we found that for targeting shorter segments, the RFdiffusion generated designs with the best metrics often made extensive beta strand interactions to targets adopting beta strand conformations. To increase the efficiency of generating such designs, we incorporated into the RFdiffusion sequence input approach the ability to define the secondary structure of the target (See Methods), to enable the specification of either the entire or a portion of the target sequence in helical, strand, or loop conformation. This is particularly important for strand conformations which can vary considerably in actual 3D coordinates; the coordinate specifying approach used by Vasquez et al^11^ for helical peptides would be less efficient for targeting strands as many trajectories would have to be carried out for beta strand conformations with different twists, etc. To explore the power of this approach, we used it to design binders to three IDR containing targets.

G3BP1 is a central node within the core stress granule (SG) network^27^ and plays a crucial role in RNA metabolism and stress response, with a disordered RNA-binding domain (abbreviated as RBD; KPGFGVGRGLAPR, 13 amino acid) mediating interactions with RNA molecules, regulating RNA metabolism, and contributing to the assembly and disassembly of stress granules. A first round of 10,000 RFdiffusion trajectories with sequence only specification of the RBD domain of G3BP1, abbreviated as G3bp1^RBD^ yielded designs with the peptide adopting a roughly 5.7 :3.8 :0.5 ratio for helix:strand:loop, respectively (Supplementary Fig. 4a), but only the 23 strand containing designs had AF2 pae_interaction < 10 and plddt_binder = 90 (Supplementary Fig. 4a-b). Based on these observations, we specified the secondary structure as a strand and conducted 10,000 trajectories. The resulting ratio of G3bp1^RBD^ conformation in the complex was 0.54:8.9:0.6 for helix:strand:loop, respectively, with 1,192 designs meeting the same filtering criteria, a ∼51 fold improvement; in all passing designs the target had a strand conformation. We narrowed these down to 78 designs by filtering on structure prediction and Rosetta interaction metrics (monomer plddt, hbonds_count, monomer RMSD, sap_score, ddg, and contact_molecular_surface). The 78 designs were subsequently expressed in E. coli and subjected to initial screening using BLI. 5 out of 78 designs were found to bind to G3bp1^RBD^, with the tightest exhibiting a binding affinity at 18 nM. Through two-side partial diffusion, we further optimized 4 of the binders (G3bp1-4, G3bp1-45, G3bp1-53, and G3bp1-77; Supplementary Fig. 4c); 40 of the 95 refined designs bound G3bp1^RBD^, with the tightest G3bp1-11 having an affinity of 11nM.

We next sought to make binders of the prion protein which is primarily found in neuronal cells in mammals. Aggregated forms of this protein are linked to prion diseases, a group of transmissible neurodegenerative disorders^28,29^. The pathological hallmark of prion diseases is the conformational conversion of the native, monomeric cellular prion protein (PrP^C^) into a misfolded and aggregated form (PrP^Sc^) characterized by a cross-β structure^30–33^. To target the amyloid core region of the prion protein, we targeted the amino acid sequence VNITIKQH (positions 180-187), specifying its secondary structure as a β-strand and conducted 20,000 trajectories. Using in silico filtering strategies similar to those employed for G3bp1^RBD^, we selected 48 designs for further validation via BLI. Among these, the tightest binder, PRI28, had a binding affinity of 14 nM (Fig. 2h) with high stability up to 95 °C (Supplementary Fig. 5a), higher affinity and specificity than generally achieved with our earlier Rosetta based β-strand targeting method^13^ (Supplementary Fig. 5b). Moreover, we found that specifying the secondary structure of the target region as a β-strand resulted in binders with higher affinity than using the target sequence information alone (14 nM from secondary structure specification (PRI28) vs 1.88 μM sequence input (PRI22), Fig. 2h and Supplementary Fig. 5c-d). after refinement through two-sided partial diffusion, the affinity of PRI22 improved to 80 nM, still weaker than PRI28 (Supplementary Fig. 5c-d).

Signal transduction via cell surface receptors is mediated by their intracellular domains, which contain long disordered regions^34,35^. Developing binders targeted at these domains would be broadly useful for co-localization imaging applications and for the modulation of receptor activation. The common cytokine receptor γ chain (common gamma chain, IL2RG) is a receptor subunit shared among the interleukin (IL) receptors for IL-2, IL-4, IL-7, IL-9, IL-15 and IL-21. Each receptor within the γc family uniquely contributes to the adaptive immune system, influencing the development of T, B, natural killer, and innate lymphoid cells^36^. To target the intracellular domain of IL2RG, we selected the amino acid sequence ERLCLVSEIP (positions 327-336) as the target region, specifying its secondary structure as a strand and conducted 40,000 trajectories. Employing *in silico* filtering strategies similar to those used for G3bp1, we selected 94 designs for further validation via BLI. Among these, one design had a binding affinity of 493 nM. Through two sided partial diffusion, we increased the binding affinity to 97 nM, and we named it IL2RG-30 (Fig. 2i); this optimized design again had high thermal stability (Supplementary Fig. 4e).

## Structure analysis of designed complexes

We obtained crystal structures of Amylin-22αβL and G3bp1-11 in complexes with their target at 1.8-Å-resolution and 2.4-Å-resolution, respectively. For Amylin-22αβL, the designed conformation comprises a helix, a strand, and an unstructured loop (Fig. 3a, left). The Amylin helix is embedded within a groove formed by the helix and strand segments of the binder. Adjacent to this, the Amylin strand pairs with a corresponding strand of the binder. The Amylin loop is predicted to be disordered based on the low per-residue AF2 pLDDT (predicted Local Distance Difference Test) (Fig. 3a, left, Supplementary Fig. 1c)^22,37^. In the crystal structure, the main helix and strand are well resolved, and closely match the computational model; the disordered loop is as anticipated not resolved (Fig. 3a-b). The Ca RMSD between the design model and the crystal structure over the backbone of the binder alone, and over the backbone of the full complex excluding the missing loop of Amylin, are 0.96 and 2.04, respectively. The backbone and sidechains at the designed binder-target interface are also in close agreement between crystal structure and design model (Fig. 3b, interface Ca and sidechain RMSD are 1.33 and 1.87, respectively).

**Figure. 3.**
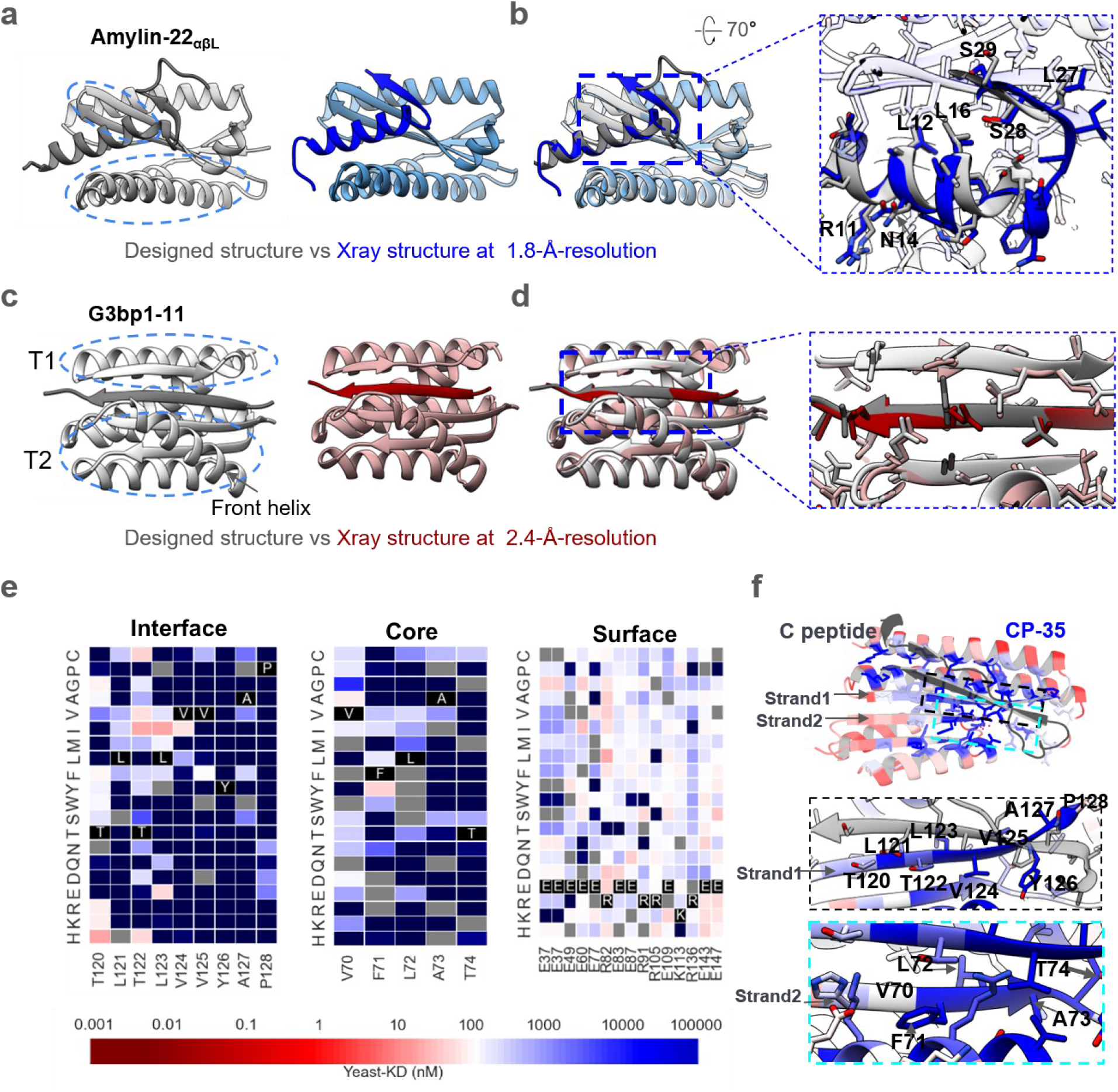
Structural characterizations. **a,** Left, the designed model of Amylin-22αβL, with target and binder proteins rendered in dim gray and gray, respectively. The helical and strand segments that create the groove in the binder, docking the helical segment of Amylin, are highlighted with blue dashed ellipsoid. Right, the crystal structure of Amylin-22αβL at 1.8 Å-resolution, with target and binder proteins rendered in blue and cornflower blue, respectively. **b,** Left, the overlay of the design model and the crystal structure of Amylin-22αβL. Right, magnified views of the regions indicated with black dotted frames in the left panel are provided to illustrate the detailed interface view of the design and crystal structure. The binder proteins are rendered with 90% transparency to enhance the visibility of the peptide target. The key residues on the Amylin are labeled to illustrate the good alignment of the key residues between designed protein and crystal structure. **c,** Left, the designed model of G3bp1-11, with target and binder proteins rendered in dim gray and gray, respectively. The two α/β topologies (T1 and T2) of the binders, forming the cleft where the target strand is positioned, are highlighted with blue dashed ellipses. The front helix of T2 is denoted by a black arrow. Right, the crystal structure of G3bp1-11 at 2.4 Å-resolution, with target and binder proteins rendered in dark red and rosy brown, respectively. **d,** Left, the overlay of the design model and the crystal structure of G3bp1-11. Right, magnified views of the regions indicated with black dotted frames in the left panel. The front helix of T2 has been surface capped to reveal the strand pairing interface. **e,** Heat maps representing C peptide-binding Kd (nM) values for single mutations in the designed interface (left), core (middle) and the surface (right). Substitutions that are heavily depleted are shown in blue, and beneficial mutations are shown in red, gray color indicates the lost yeast strains. For the interface region, we highlighted and showcased strand 1 (indicated by the arrow), which serves as the primary interaction secondary structure with the C peptide. For the core region, we showcased the right segment of strand 2 (indicated by the arrow), representing a main core region that does not form interactions with the C peptide. For the surface region, we selected the most exposed surface residues that don’t form any connections with other residues (Supplementary Fig. 6c). Full SSM map over all positions for CP35 is provided in Supplementary Fig. 6b. **f,** Top, designed binding proteins are colored by positional Shannon entropy from site saturation mutagenesis, with blue indicating positions of low entropy (conserved) and red those of high entropy (not conserved). Bottom, zoomed-in views of central regions of the design interface and core with the C peptide.

In the G3bp1-11 design model, the peptide is in a β-strand conformation and lies within a cleft formed by two α/β structures, T1 and T2, in the designed binder, pairing with two adjacent strands (Fig. 3c). An additional helix in T2 also interacts with the target, potentially enhancing binding affinity and specificity (Fig. 3c, Supplementary Fig. 6a). The crystal structure of G3bp1-11 closely recapitulates the design model, with the peptide clamped in a β-strand conformation (Fig. 3c-d, Ca RMSD 0.8 Å for entire complex between design and crystal structure) with the interface residues nearly perfectly aligned with the design model structure (Fig. 3c-d, interface Ca and sidechain RMSD are 0.86 and 2.29, respectively).

We were unable to solve crystal structures of the CP binders, so we instead obtained a lower resolution structural footprint of the binding site by generating a site saturation mutagenesis library (SSMs) for CP-35 in which every residue was substituted with each of the 20 amino acids one at a time. Next generation sequencing before and after FACS sorting for CP binding revealed that residues at the binding interface and protein core were largely conserved (Fig. 3e-f and Supplementary Fig. 6b-c), supporting the design model.

## Specificity of designed binders

We investigated the specificity of the binders by carrying out all by all binding experiments (Fig. 4). BLI binding characterization of 9 binders against 6 targets showed that the designs had high specificity for their intended peptide targets. Very weak off target binding was observed at high concentrations in two cases: VP48 weakly bound Amylin above 800 nM, perhaps reflecting the ∼50% helical content of both peptides (specificity could potentially be further improved through another round of partial diffusion, or decreasing the helical percentage through secondary structure specification) and G3BP1-11 weakly bound IL2RG at 2 uM. Overall, the much higher on-target than off-target binding suggests the binders should be broadly usable as affinity reagents.

**Figure. 4.**
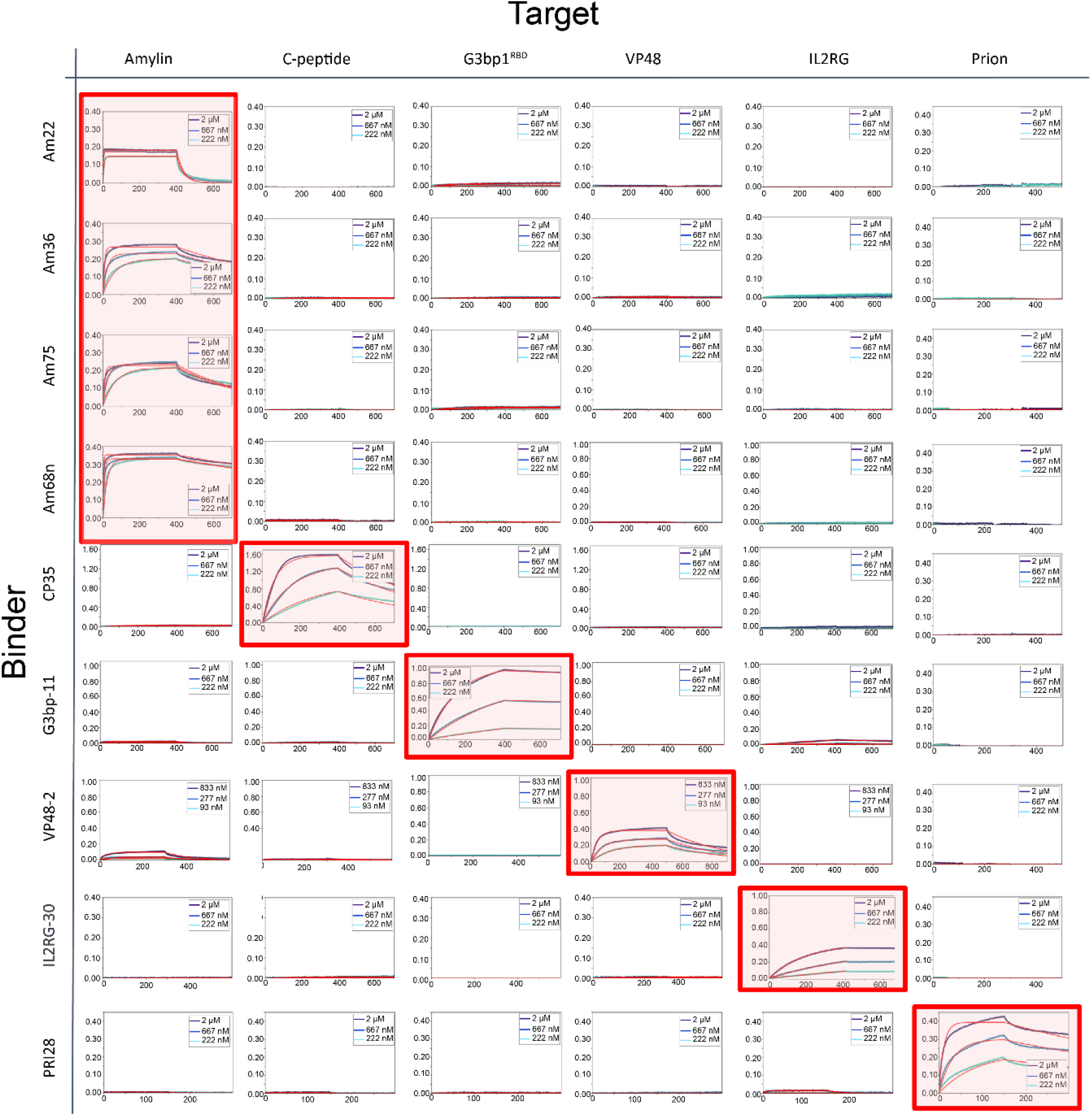
Specificity profile of designed binders in BLI. Biotinylated peptides were immobilized onto octet streptavidin biosensors at equal densities and incubated with all binders in separate experiments at three concentrations (2, 0.667 and 0.222 μM except VP48 binder at 0.833, 0.277 and 0.093 μM). Amylin-68nαβ, Amylin-36αβ, Amylin- 75αα, Amylin-22αβL are abbreviated as Am68n, Am36, Am75 and Am22, respectively. The designed on-target interactions are indicated with a light red background.

### Designed binders colocalize with their targets in mammalian cells

To examine whether the designs could fold properly and bind to the target proteins in mammalian cells, we knocked out the endogenous IL2RG in HeLa cells using CRISPR-Cas9, and then transfected the cells with a construct encoding IL2RG fused to EGFP. When cells were additionally transfected with mScarlet-labeled IL2RG binder IL2RG-30, colocalization of GFP and mScarlet was observed, indicating binding (Fig. 5a). In IL2RG knockout cells transfected only with IL2RG-30-mScarlet, no colocalization was observed (Fig. 5a, left), confirming that the interaction occurs through the designed interface.

**Figure. 5.**
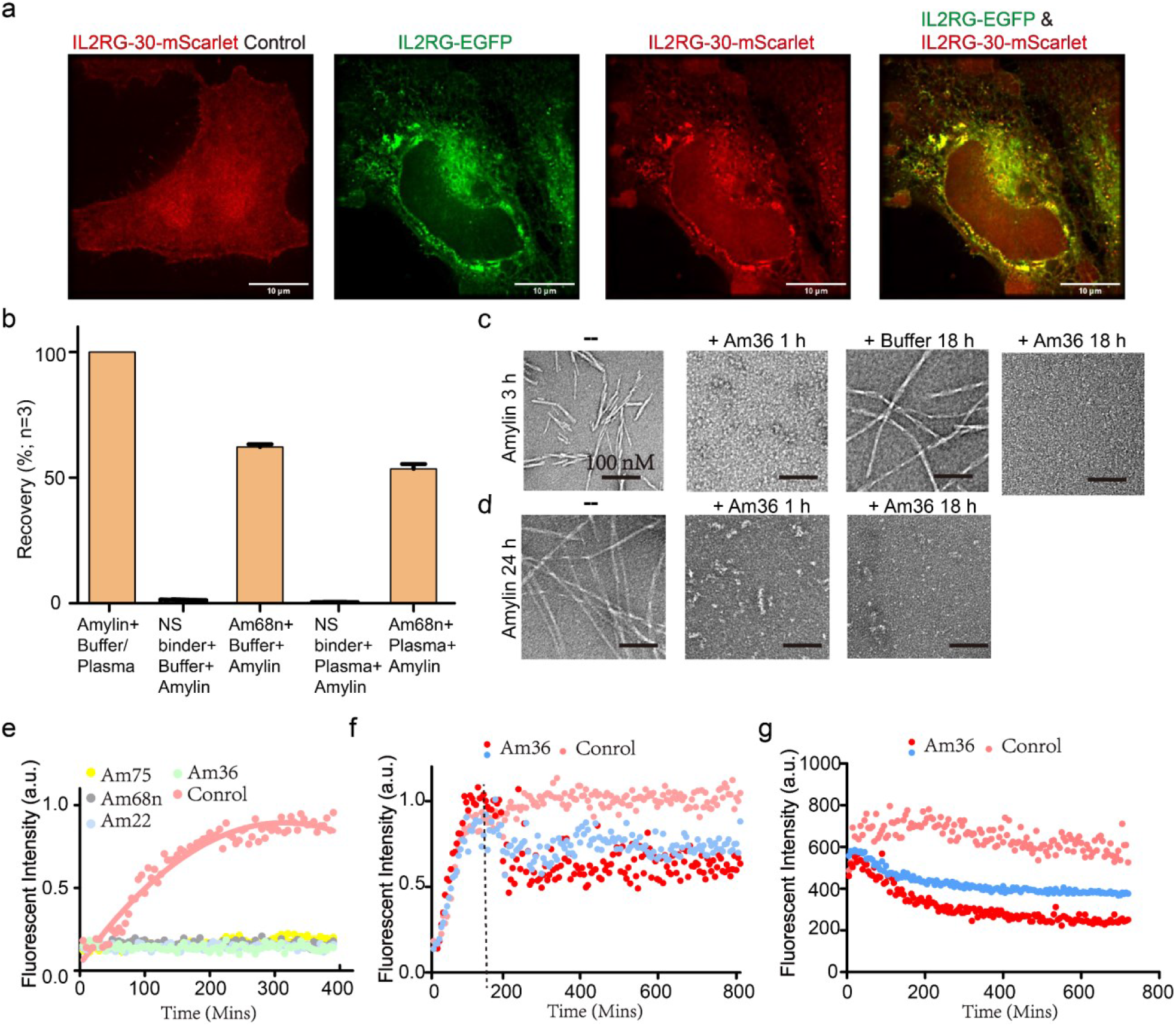
Applications of designed binders. **a,** Colocalization of binder IL2RG-30 and target membrane receptor IL2RG in HeLa Cells. Cells with endogenous IL2RG knocked out express only the red fluorescent mScarlet-tagged binder IL2RG-30, which is uniformly distributed throughout the cell (left). In contrast, cells co-expressing green EGFP-tagged IL2RG and red mScarlet-tagged IL2RG-30 show specific colocalization of both proteins. **b**, The LC–MS/MS recovery percent of Amylin from PBS-0.1% CHAPS buffer and EDTA-anticoagulated plasma was compared between BSA-blocked tosyl-activated bead, an off- target binder, and amylin-targeted binders (Am68n). Percent recovery was calculated using the peak area of a sample of pure amylin peptide in elution solvent as the denominator (i.e., 100% recovery of the peptide). Error bars represent SD (n=3). **c-d**, Visualization of fibril dissociation by Amylin-36αβ binder using negative staining electron microscopy. panels (c) and (d) demonstrate the dissociation of existing fibrils at elongation phase (c) and mature phase (d) following the addition of Amylin-36αβ. Scale bars, 100 nM. **e**, Thioflavin T (ThT) assay revealed that all 4 binders could strongly inhibit fibril formation at molar ratio of binder to Amylin 1:4. **f**, Amylin- 36αβ could dissociate fibrils at elongation phase in concentration-dependent manner. The Tht assay was performed since the Amylin monomer, Amylin-36αβ was added at 3h when Amylin fibrils were at elongation phase, marked with a dotted line. Red dot and blue dot indicate that Amylin- 36αβ to Amylin is 1:4 and 1:40, respectively. **g**, Tht assay was performed after the mature Amylin fibrils were formed for 24 h, at the same time, Amylin-36αβ was added, the data revealed that fibril fluorescence decreased in a concentration-dependent manner. Red dot and blue dot indicate that Amylin-36αβ to Amylin is 1:4 and 1:40, respectively.

## Enrichment for LC–MS/MS detection

We explored the use of amylin binder Amylin-68n as a capture agent for immunoaffinity enrichment combined with liquid chromatography–tandem mass spectrometry (LC–MS/MS), a general platform for detecting low-abundance protein biomarkers in human serum^38^. We prepared Amylin-binder-conjugated beads as described in the Methods. Amylin enrichment was calculated based on detection of intact, alkylated amylin in either human plasma or simplified PBS-CHAPS matrix^39^ (Methods). We found that the designed binder enabled capture of Amylin from buffer and human plasma supplemented with Amylin (the endogenous levels are too low for reliable detection) with recoveries of 62.2% and 53.5%, respectively (Figure. 5b).

### Designs inhibit Amylin fibril formation and dissociate existing fibrils

Amylin fibril formation is implicated in type 2 diabetes, where the aggregation of amylin into insoluble fibrils contributes to islet amyloid deposition and β-cell dysfunction^40^. We investigated the effect of four binders—Amylin-68nαβ, Amylin-36αβ, Amylin-75αα and Amylin-22αβL—on Amylin fibril formation. At a binder to Amylin molar ratio of 1:4, with concentrations of 40 μM for Amylin and 10 μM for binders, all binders completely inhibited fibril formation (Fig. 5e). Further tests with Amylin-22αβL and Amylin-36αβ at binder to Amylin molar ratios of 1:4, 1:40, and 1:400 revealed a concentration-dependent retardation of fibril formation (Supplementary Fig. 7a). Inhibition of fibril formation was also observed by negative stain electron microscopy (NS- EM), with Amylin-22αβL and Amylin-36αβ at binder to Amylin molar ratios of 1:4. Addition of Amylin-36αβ blocked fiber formation at both 1 h and 18 h, whereas some short fibrils were observed 18 hours post-addition of Amylin-22αβL (Supplementary Fig. 7b-c).

We next investigated whether the amylin binders were able to disaggregate pre-formed amylin fibrils. We generated short Amylin fibrils by incubating the peptide at 40 μM for 3 hours at 37 °C, to reach the elongation phase, and then incubated with 10 μM Amylin-36αβ. NS-EM revealed no fibrillar structures after treatment with Amylin-36αβ at both 1 h and 18 h time points (Fig. 5c). Thioflavin T (ThT) assays with Amylin-36αβ added at the 3-hour Amylin fiber stage also showed fiber disassembly in a design concentration-dependent manner (Fig. 5f).

To test whether Amylin-36αβ could dissociate mature fibrils that had formed over 24 hours at 10 μM, we incubated them with 10 μM of the binder. Small oligomers were still observed at 1 hour, but were completely dissociated by 18 hours (Fig. 5d). Fibril ThT fluorescence again decreased in a designed binder concentration-dependent manner (Fig. 5g).

## Discussion

Our results demonstrate the utility of RFdiffusion in designing binders for IDPs ranging from 30- 40 amino acids in length in diverse conformations, expanding its applicability beyond helical peptides. The ability to target IDPs without specifying the target structure is important as such proteins have no single defined conformation. During the design process, the target protein samples a wide range of possible conformations as the designed binding protein diffuses around it; the co-folding of design and target effectively enables the selection of conformations particularly suitable for binding. The versatility of our approach is highlighted by the design binders for Amylin in diverse conformations while consistently forming the Amylin peptide disulfide.

For shorter peptides which can adopt beta strand like conformations, we show the introduction of a secondary structure type specification feature within the RFdiffusion model enables targeting of peptides in the beta strand conformation. The generated structures resemble previous strand targeting designs generated using Rosetta, but exhibit higher specificity and binding affinity.

The binders and approaches described here could be broadly useful given the current difficulty in targeting IDPs and IDRs, and the important roles these play in both normal physiology and disease. For example, the Amylin binder both inhibits the formation of Amylin fibers and dissociating pre-existing fibers, which could have therapeutic utility. Additionally, it facilitates the enrichment and detection of Amylin using mass spectrometry. The designed binders bind their targets in cells, as illustrated by the colocalization of PRI28 with the intracellular tail of the IL2 receptor gamma subunit, opening up new ways of modulating cytokine signaling in feedback loops for adoptive cell therapies and other applications.

## Supplementary data

**Supplementary figure. 1.**
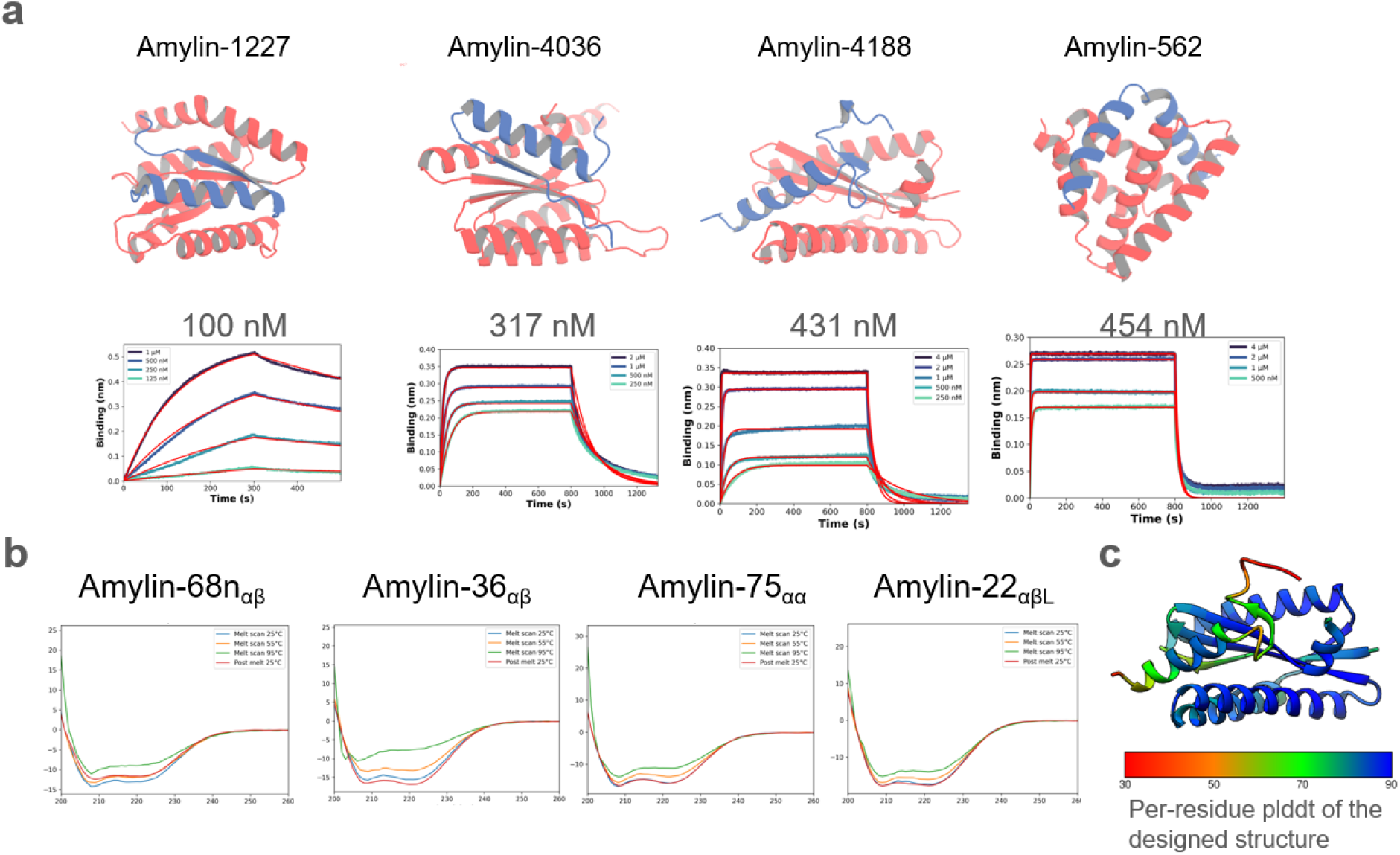
Diffusing de novo peptide binder design to Amylin. **a**, Top, the designed structures of four initial hits, Amylin-1227, -4036, -4188, -562, which serve as starting point of two sided partial diffusion. Bottom, the BLI result of the four hits revealing the binding affinity of the 4 initial hits are 100, 317, 431, 454 nM, respectively. **b,** Circular dichroism data show that the optimized binders have helical secondary structure and is stable up to 95 °C (inset). **c,** The per residue pLDDT (predicted Local Distance Difference Test) plotting of Amylin- Am22 complex in design.

**Supplementary figure. 2.**
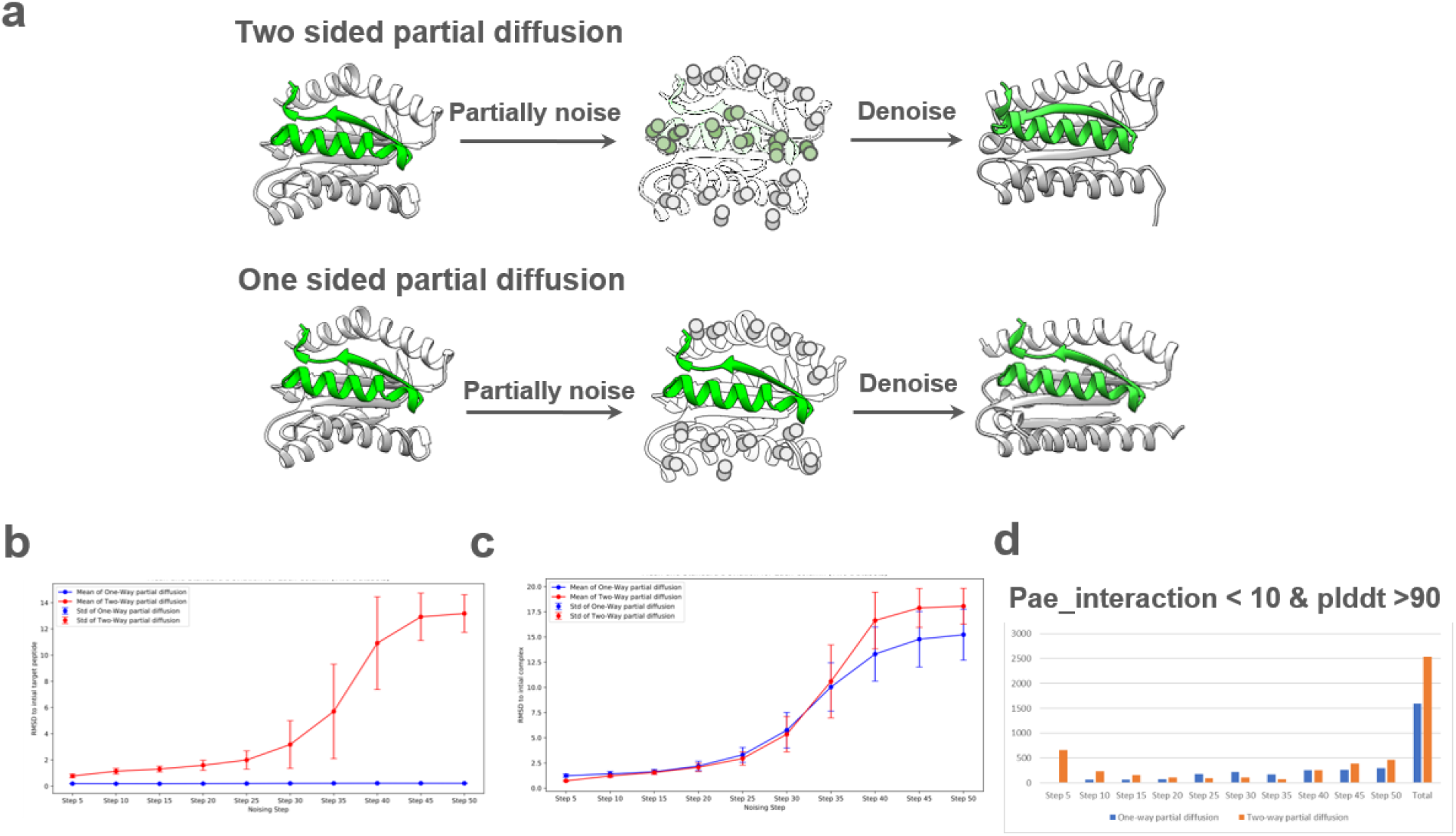
Two sided partial diffusion and the comparison with one sided partial diffusion. **a,** Top, two sided partial diffusion allows simultaneous conformational changes in both the target and the binder. Bottom, one sided partial diffusion solely diversifies the conformation of the binder while keeping the target fixed. **b,** Two sided partial diffusion (in red) diversifies the target while one sided partial diffusion (in blue) keeps the target fixed. **c,** The peptide-binder complex diverse magnitudes of two sided (in red) and one sided partial diffusion (in blue) remain comparable before nosing step 35, after step 35, the diverse magnitude of two sided parietal diffusion is larger than one sided one. **d,** Take the interface pAE <10, pLDDT >90 as cutoff criteron, two sided partial diffusion yielded designs with generally better metrics than one sided diffusion. At steps 25, 30, and 35 exclusively, one-sided partial diffusion exhibited superior performance. However, in practical cases, we typically operate within fewer than 25 steps to remain the main features of parent structure.

**Supplementary figure. 3.**
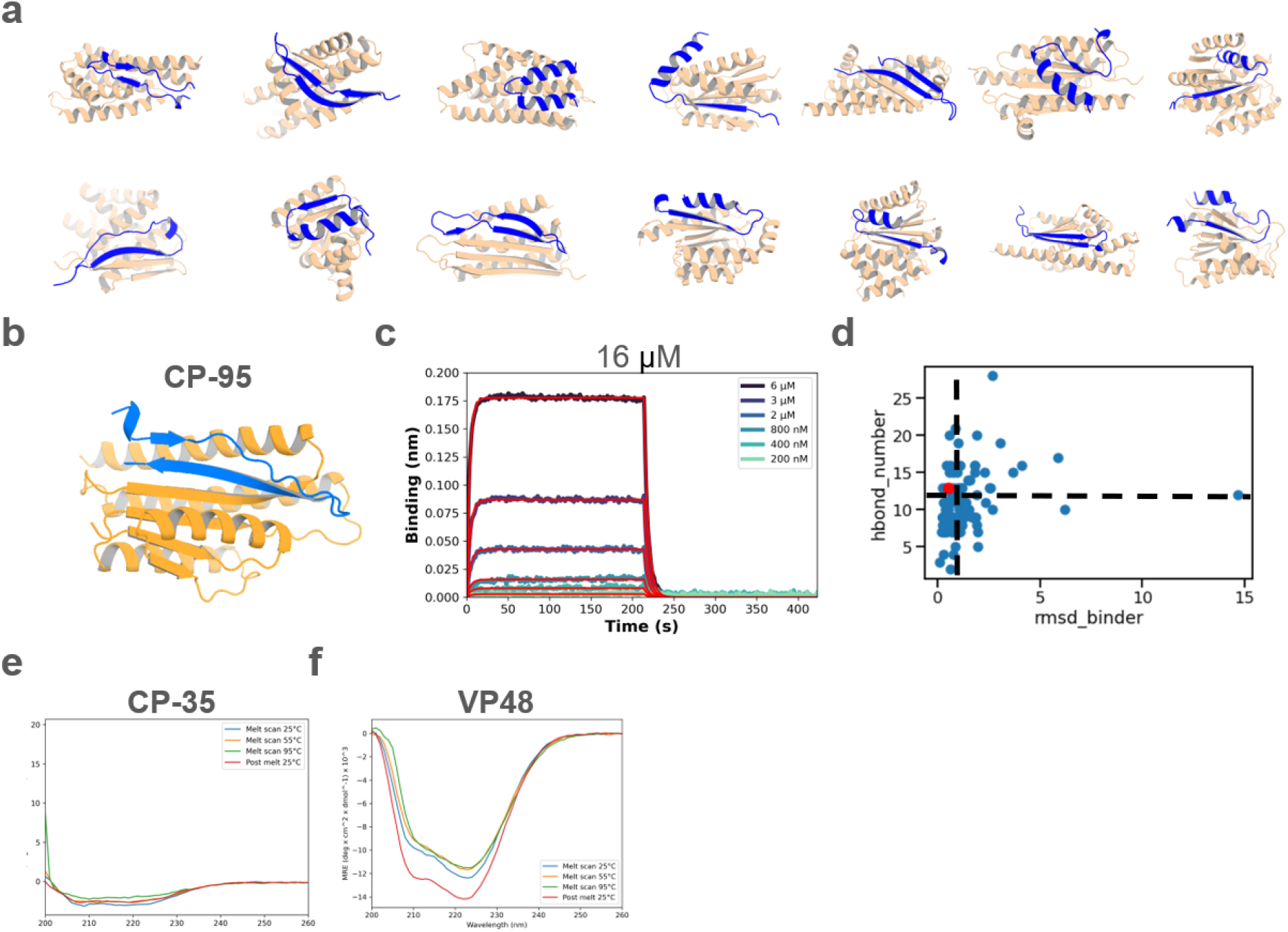
Diffusing de novo peptide binder design to C peptide. **a,** Sequence-input diffusion was carried out, allowing C peptide to sample diverse conformations. The diverse conformations of C peptide and protein binder are rendered in blue and wheat color, respectively. **b,** Design model of the initial hit CP-95 which was also the starting point of two-sided partial diffusion. **c**, the BLI data revealed the binding affinity of the initial hit is 16 μm. **d,** Scatter plot showing the distribution of designs based on the number of hydrogen bonds (hbond_number) and the RMSD of the binder (rmsd_binder). Each blue dot represents a design, while the red dot marks a validated hit. The dashed black lines indicate the cutoff values based on the initial hit criteria (hbond_number = 13 and rmsd_binder = 0.545). Analysis revealed that only 6 out of 96 designs met these criteria (hbond_number > 13 and rmsd_binder < 0.545), indicating a low success rate. **e-f,** Circular dichroism data show that the binder CP35 (e) and VP48 (f) have helical secondary structure and is stable up to 95 °C (inset).

**Supplementary figure. 4.**
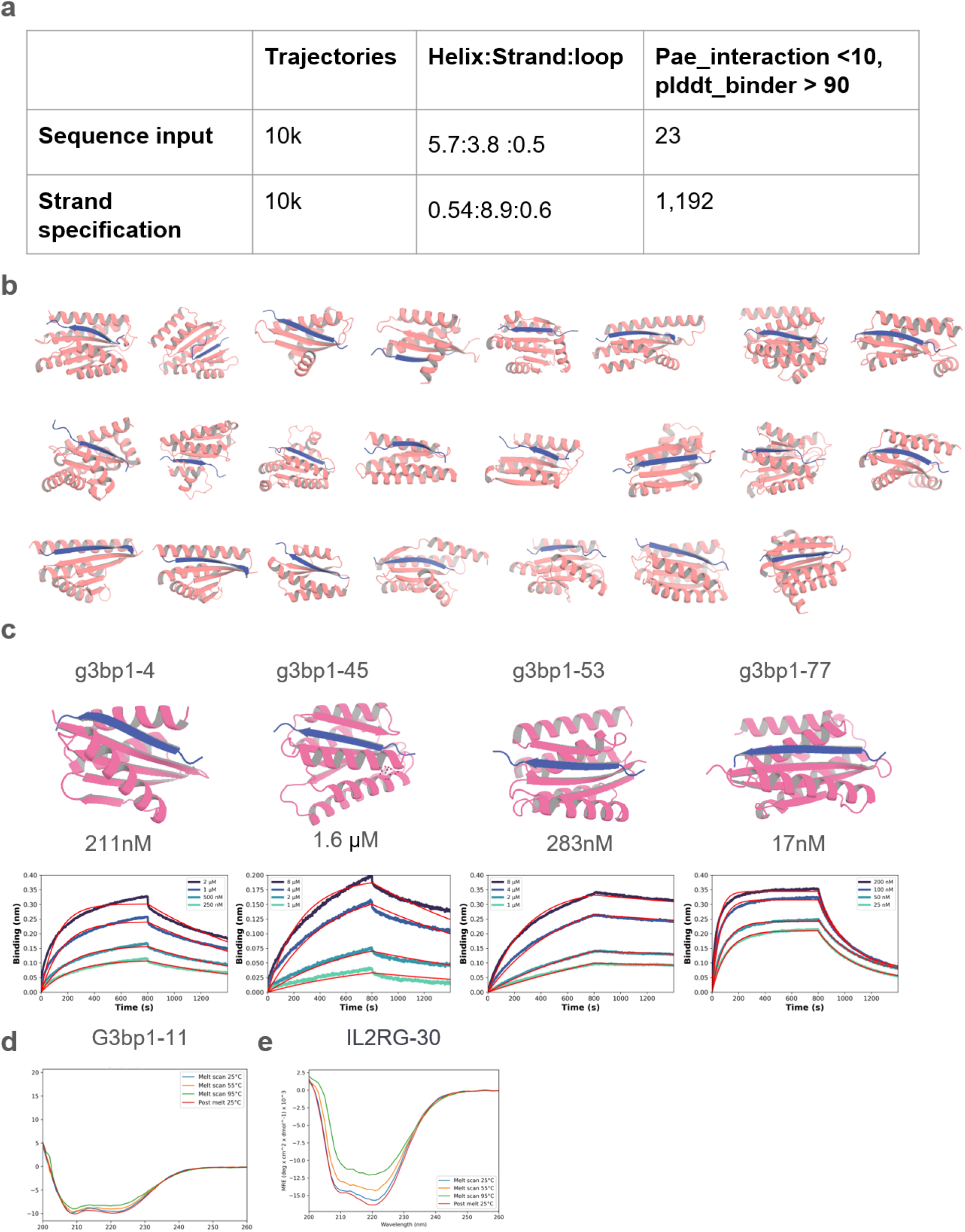
Diffusing de novo peptide binder design to G3BP1^RBD^. **a,** Comparative analysis of structural outcomes between sequence input and strand specification approaches in protein design. The table presents the number of trajectories (10k) and the distribution of secondary structures (Helix:Strand: Loop) for both methods. This table counts the successful cases where the Pae_interaction < 10 and the plddt_binder score = 90, noting 23 successes with sequence input and 1,192 with strand specification. This reflects an approximately 51-fold increase in efficacy with the strand specification method, highlighting its superior performance in achieving desired structural configurations. **b**,The 23 successful cases designed using sequence input RFdiffusion all feature targets in strand conformation. **c**, Design models and BLI data of the 4 initial hits of G3BP1^RBD^ which was also the starting point of two sided partial diffusion. **d and e,** Circular dichroism data show that the G3bp1-11 binder (d) and IL2RG-30 binder (e) have helical secondary structure and are stable up to 95 °C (inset).

**Supplementary figure. 5.**
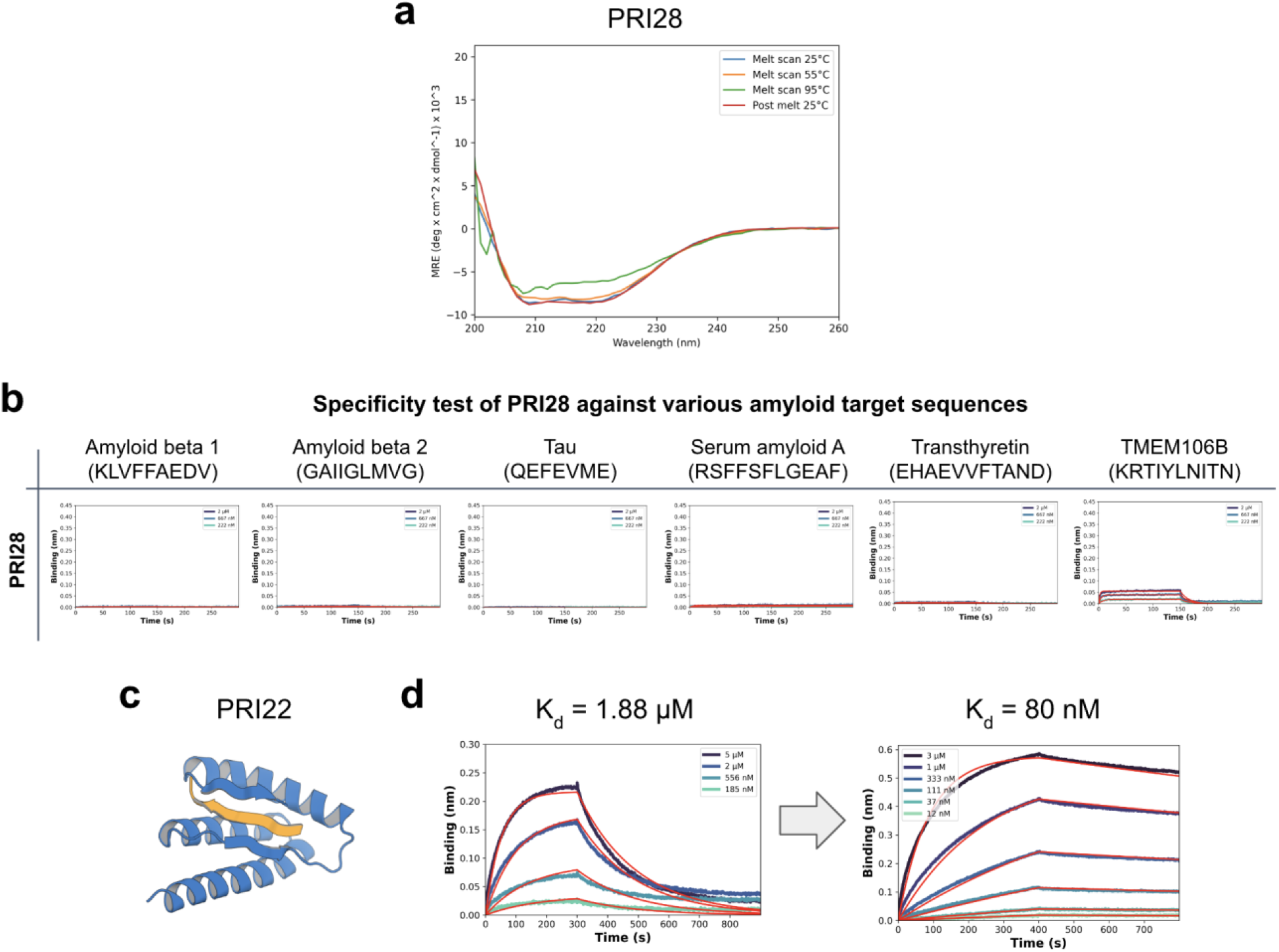
Diffusing de novo peptide binder design to prion protein. **a**, Circular dichroism data show that the PRI28 binder has helical secondary structure and is stable up to 95 °C. **b**,The specificity test for prion binder PRI28 against various amyloid target sequences showed that PRI28 is highly specific, with some cross-reactivity observed only with TEME106B, related to Fig. 2h. **c,** The design model of PRI22, designed using target sequence information alone, is shown. **d**, The BLI data revealed that the binding affinity of PRI22 is 1.88 μM (left), which improved to 80 nM after two-sided partial diffusion (right).

**Supplementary figure. 6.**
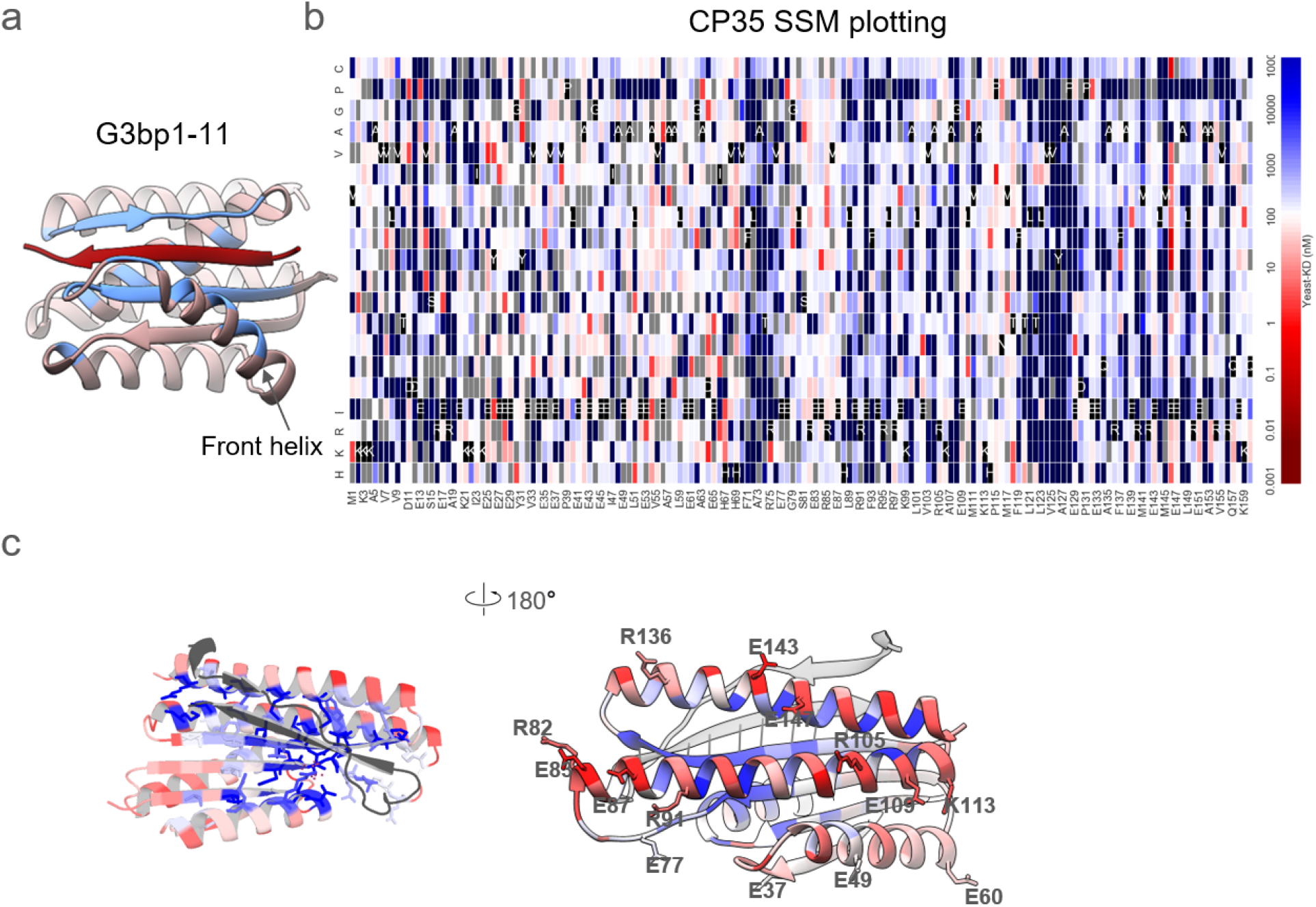
SSM analysis of CP35. **a,** The crystal structure of G3bp1-11, positioned 4 Å away from the target on the binder, is marked in blue. **b**, Full SSM maps for the design of CP35. **c**, Zoomed-in views of the residues presented in the surface region, as shown in Figure 3e.

**Supplementary figure. 7.**
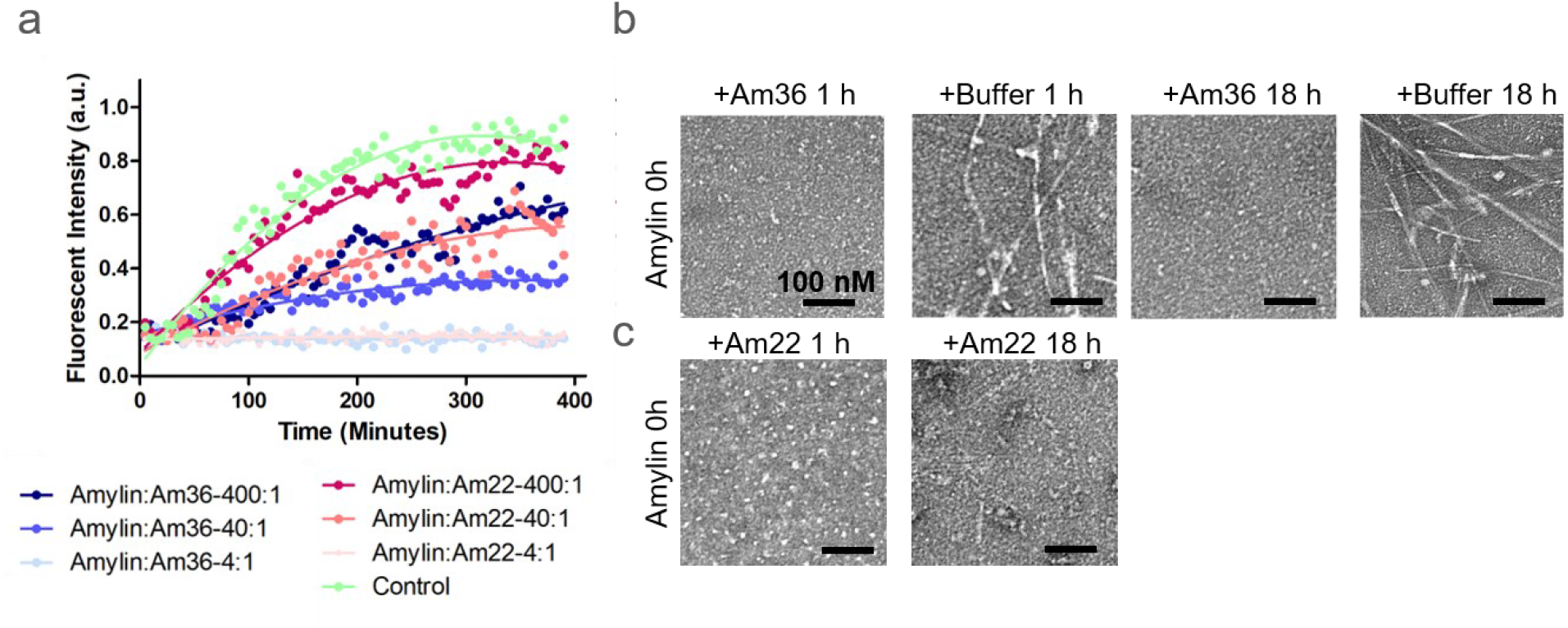
Designs inhibit Amylin fibril formation and dissociate existing fibrils a,Amylin binders Amylin-22αβL and Amylin-36αβ inhibit fibril formation in a concentration-dependent manner. The initial concentration of Amylin monomer was 10 μM, with subsequent additions of binders at 2.5 μM, 0.25 μM, and 0.025 μM, establishing molar ratios of binder to Amylin of 1:4, 1:40, and 1:400, respectively. b-c, Negative stain electron microscopy images were taken of 40 μM Amylin monomer samples following the addition of 10 μM Amylin-36αβ (b) and Amylin-22αβL (c) at 1 hour and 18 hours, respectively. Scale bars, 100 nM.

**Supplementary table 1.**
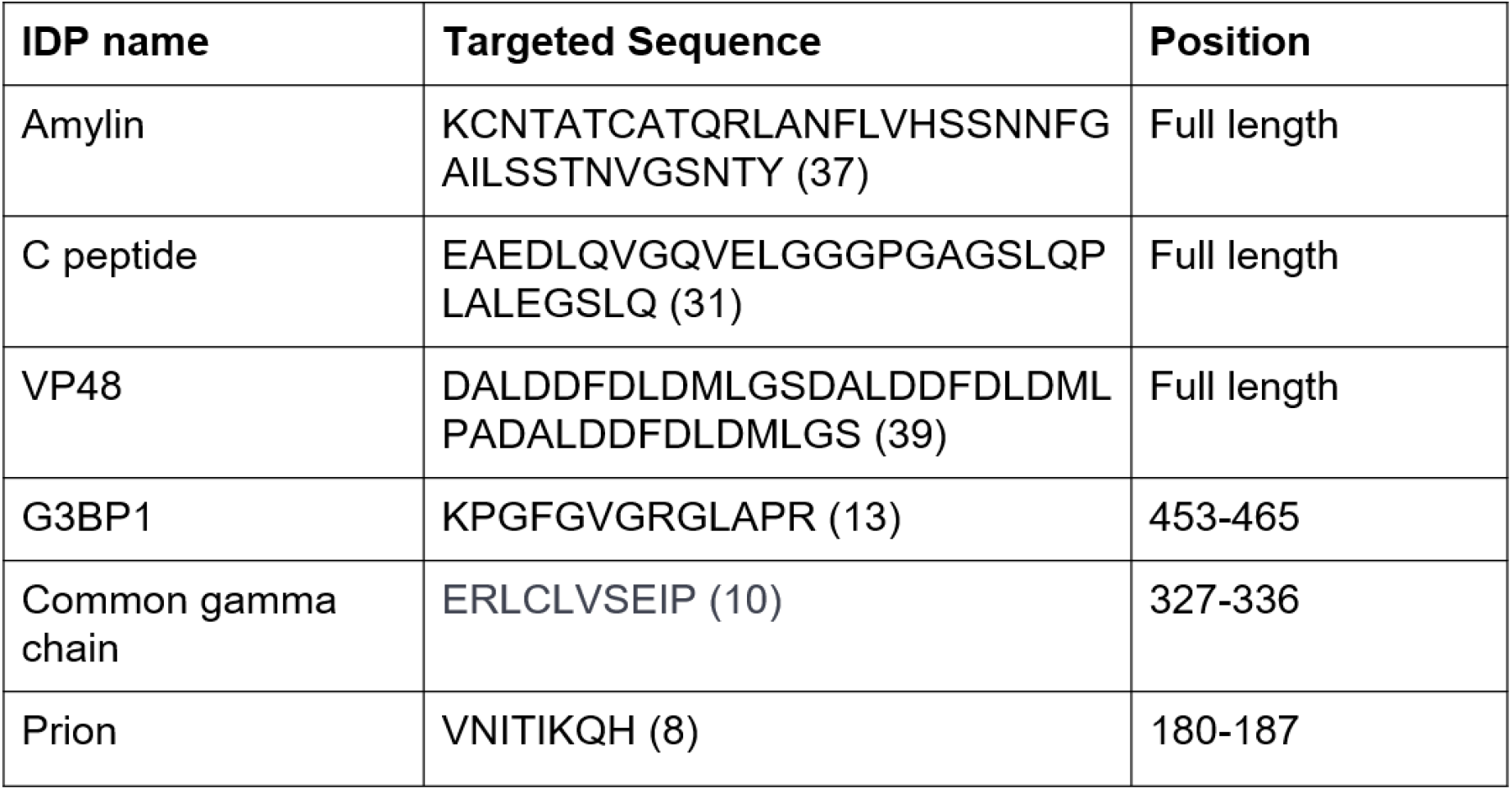
Summary of IDPs and IDRs in the study, detailing each protein’s sequence and positional data within their respective structures.

**Supplementary Table 2.**
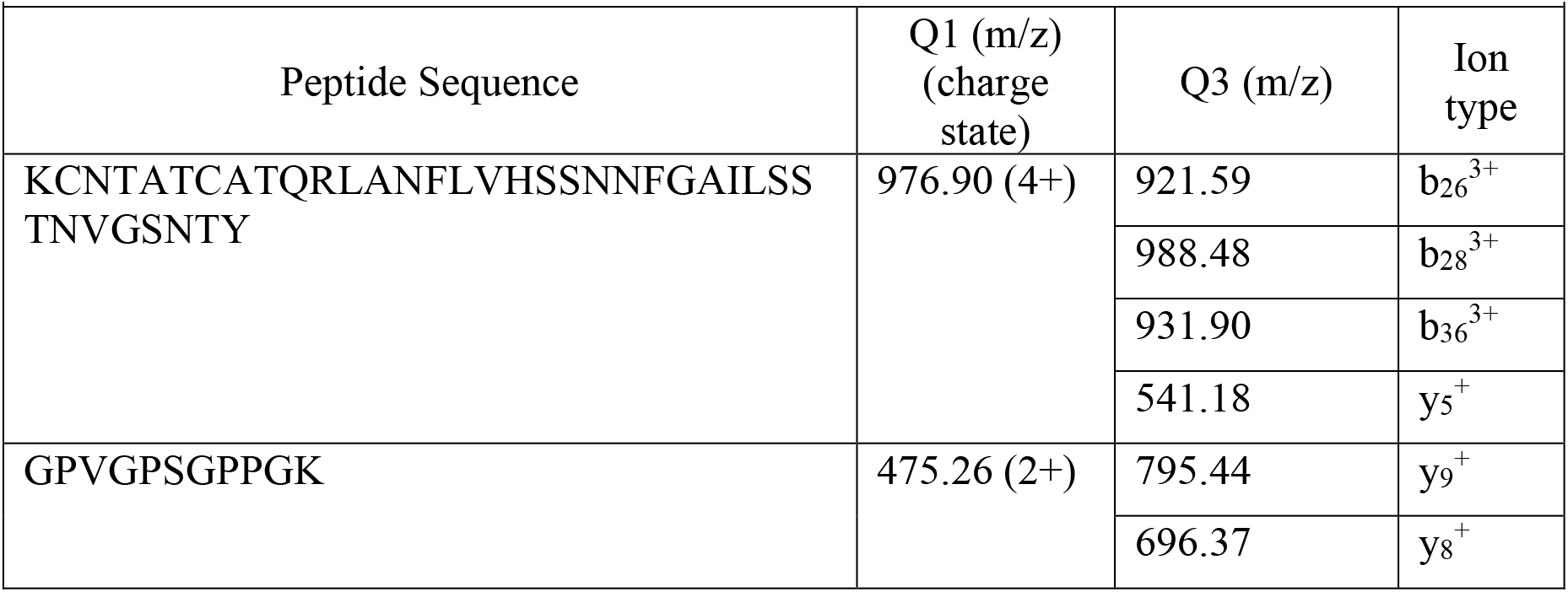
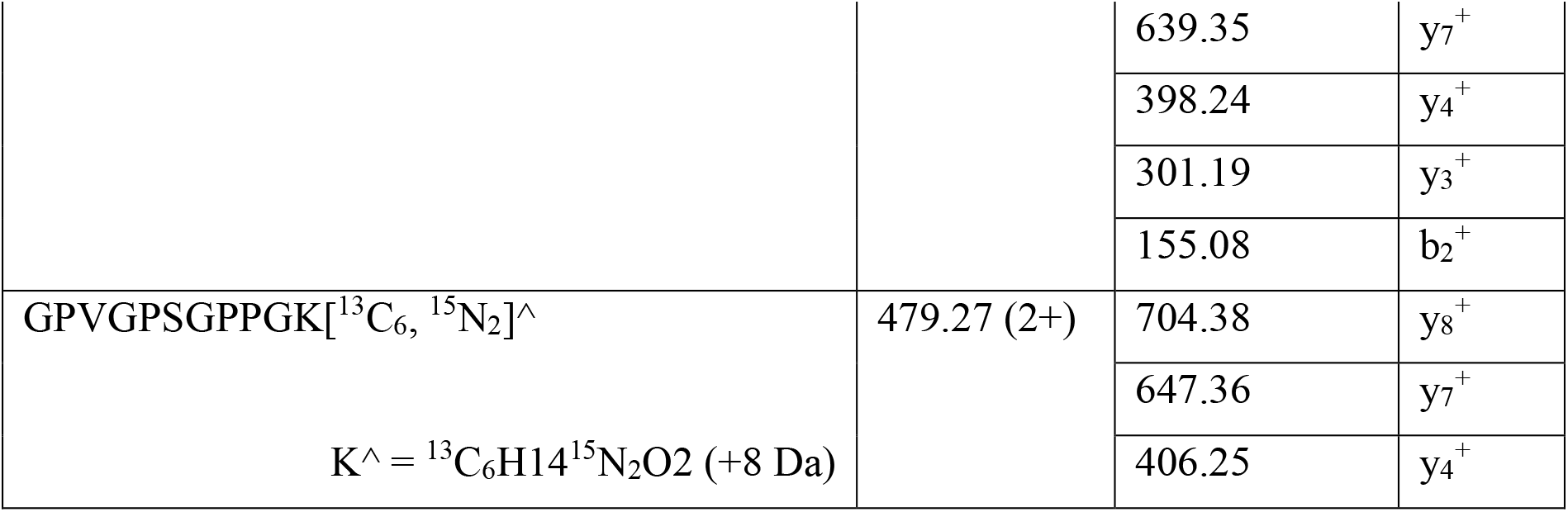
Amylin transitions monitored

**Supplementary Table3.** Liquid chromatography parameters

| Supplementary Table3. Liquid chromatography parameters |  |  |
| --- | --- | --- |
| Mobile phase | Phase A: 0.2% formic acid in water |  |
|  | Phase B: 0.2% formic acid in acetonitrile |  |
| Column | Acquity UPLC HSS T3 1.8μm (C18, 2.1x50 mm, pore size 100 Å) (Waters, Milford, MA, P/N 186003539) |  |
| Temperature | 45°C |  |
| Flow rate | 0.3 mL/min |  |
| Injection volume | 10μL |  |
| Gradient | 0-0.5 min | 20% B at 0.3 mL/min |
|  | 7.5 min | 60% B at 0.3 mL/min |
|  | 9.5 min | 98% B at 0.3 mL/min |
|  | 11.0 min | 98% B at 0.3 mL/min |
|  | 11.1 min | 20% B at 0.3 mL/min |
|  | 12.5 min | 20% B at 0.3 mL/min |

**Supplementary Table 4.**
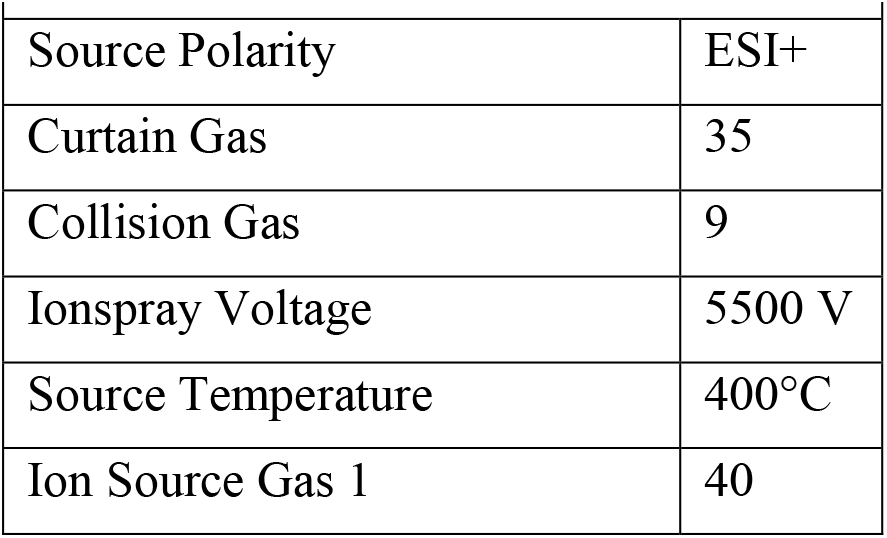
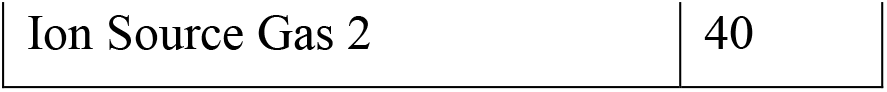
Mass spectrometry parameters

**Supplementary table 5.** Data collection and refinement statistics.

|  | Amylin-22<sub>αβ</sub>L (PDB Code: 9CC5) | G3bp1-11 (PDB Code: 9CC6) |
| --- | --- | --- |
| Resolution range | 33.32 - 1.87 (1.94 - 1.87) | 31.43 - 2.4 (2.48 - 2.4) |
| Space group | *P* 2<sub>1</sub> 2<sub>1</sub> 2<sub>1</sub> | *P* 2<sub>1</sub> |
| Unit cell | 33.33, 34.51, 127.68; 90, 90, 90 | 38.43, 42.39, 40.63; 90, 105.53, 90 |
| Unique reflections | 12855 (1372) | 4698 (410) |
| Multiplicity | 6.6 (6.8) | 5.3 (5.0) |
| Completeness (%) | 99.38 (99.28) | 93.38 (92.0) |
| Mean I/sigma(I) | 7.94 (1.22) | 11.0 (2.2) |
| Wilson B-factor | 28.60 | 38.45 |
| R-merge | 0.1598 (1.79) | 0.033 (0.623) |
| R-pim | 0.06738 (0.7403) | 0.032 (0.152) |
| CC<sub>1/2</sub> | 0.999 (0.769) | 0.993 (0.956) |
| Reflections used in refinement | 12756 (1372) | 4697 (410) |
| R-work | 0.2256 (0.2949) | 0.2199 (0.2709) |
| R-free | 0.2721 (0.3400) | 0.2567 (0.2856) |
| Number of non-hydrogen atoms | 1335 | 1197 |
| macromolecules | 1289 | 1190 |
| solvent | 46 | 7 |
| Protein residues | 163 | 150 |
| RMS(bonds) | 0.015 | 0.003 |
| RMS(angles) | 1.31 | 0.60 |
| Ramachandran favored (%) | 95.60 | 98.63 |
| Ramachandran allowed (%) | 4.40 | 1.37 |
| Ramachandran outliers (%) | 0.00 | 0.00 |
| Average B-factor | 35 | 46 |
| macromolecules | 35 | 46 |
| solvent | 40 | 43 |

Supplementary Video Legends

Supplementary Video 1

A video of the sequence input diffusion trajectory for the fully diffused Amylin-binder complex.

## Methods

### De novo peptide binder design given only sequence input using RFdiffusion and ProteinMPNN

For each target, approximately ten to fifty thousand diffused designs were generated given only sequence input of the target. The resulting library of backbones were sequence designed using ProteinMPNN, followed by AF2+initial guess^23^. The resulting designs were filtered based on interface pAE, pLDDT. In addition, AF2 monomer was performed using only the binder sequence without the peptide to filter based on the monomer pLDDT of the binder and RMSD to the binder design model. Subsequently, FastRelax was executed to obtain Rosetta metrics. The resulting binders were then further filtered based on criteria including contact_molecular_surface, ddG, SAP score and the numbers of hydrogen bonds. Specific filtering criteria were carefully selected to narrow down the set to 48 to 96 designs for each target.

### Two sided partial diffusion to optimize binders

Partial diffusion enables the input structure to be noised only up to a user-specified timestep instead of completing the full noising schedule. The starting point of the denoising trajectory is therefore not a random distribution. Rather, it contains information about the input distribution resulting in denoised structures that are structurally similar to the input. Unlike one sided directional partial diffusion, which solely diversifies the conformation of the binder while keeping the target fixed, two sided partial diffusion allows simultaneous conformational changes in both the target and the binder. The input designs were subjected to 15 noising timesteps out of a total of 50 timesteps in the noising schedule, and subsequently denoised. Approximately ten to fifty thousand partially diffused designs were generated for each target. The resulting library of backbones were sequence designed using ProteinMPNN, followed by AF2+initial guess^23^. The resulting designs were filtered in the same way as the designs from the aforementioned sequence input diffusion process.

### Integrating secondary structure specifications into RFdiffusion

To permit specification of the secondary structure (but not three-dimensional coordinates) of the peptide target, a modified version of RFdiffusion was trained that permits specification of the secondary structure of a region, along with its sequence. The training strategy largely followed that used to train previous RFdiffusion models ^11,14^, with some modifications. A summary is provided below.

Overview of “base” RFdiffusion Training: Rfdiffusion^14^ is a denoising diffusion probabilistic model (DDPM), which is fine-tuned from the RoseTTAFold structure prediction model^22,42^. In RFdiffusion, the N-Ca-C frame representation (translation and orientation) of protein backbones^22,43^ is used, and, over 200 discrete timesteps, these backbone frames are corrupted following a defined forward noising process that noises these frames to distributions indistinguishable from random distributions (three-dimensional Gaussian distribution for translations, and uniform SO(3) distribution for rotations). RFdiffusion is trained to reverse this noising process, predicting the true (X0) protein structure at each timestep of prediction (starting from randomly sampled translations and rotations). Successive predictions are used to “self-condition” predictions through an inference trajectory, and mean squared error (MSE) losses minimize the error between forward and reverse processes. Full details of training are described in Watson et al^14^.

Modifications to permit secondary structure specification of the target: As in the original RFdiffusion fine-tuned for protein binder design, RFdiffusion was trained 50% of the time on single chains from the Protein Data Bank (PDB) < 384 amino acids in length, and 50% on hetero- complexes. In the latter case, one chain (< 250 amino acids in length) was designated the “binder”, and when necessary the other “target” chain was radially cropped around the interface (to 384 – the length of the “binder” residues). For single chain examples, 20% of the time, the whole backbone was noised, and in the other 80% of cases 20-100% of the protein backbone was noised. For hetero-complex examples, the whole “binder” chain was noised. Additionally, and in contrast to the original RFdiffusion model trained for protein binder design, up to 50% of the noised monomer structure had sequence provided in the noised region. For hetero-complexes, up to 50% of the target chain backbone was also noised, while its sequence was provided to RFdiffusion. This permits RFdiffusion to condition on the sequence of the target chain in the absence of three-dimensional structure.

To permit specification of the secondary structure of the target (when three-dimensional coordinates are not provided), secondary structure and “block adjacency” ^14^ information were provided to RFdiffusion in exactly the manner described in Watson et al^14^. Briefly, 50% of the time, RFdiffusion was provided with a (partially masked; 0-75%) secondary structure of the example protein chain/hetero-complex, and (an independently-sampled) 50% of the time a (partially masked; 0-75%) “block adjacency” of the protein chain/hetero-complex. Additionally, 50% of the time, the whole inter-chain “block adjacency” was masked in hetero-complex examples. This permits RFdiffusion to condition on a (partially) pre-specified secondary structure (and/or adjacency information) of the target peptide. This version of RFdiffusion was trained for seven epochs.

To design binders using RFdiffusion through secondary structure specification, foreach target, approximately ten thousand diffused designs were generated through sequence input of the target with the additional secondary structure specification. The resulting library of backbones were sequence designed using ProteinMPNN^21^, followed by AF2+initial guess^23^. The resulting designs were filtered in the same way as the designs from the aforementioned sequence input diffusion process.

### Backbone extension for VP48 binder design

During the design campaign, it was noticed not all designs provided sufficient interactions to the whole sequence of the target, especially the loopy regions. To explore and guide RFdiffusion to make more interactions around certain regions, we selected 20 AF2 passing designed complexes from the round one design campaign, based on the above criteria and manual selection. For each base design, we requested RFdiffusion to extend the binder backbone with 10-20 amino acids from either N terminal, or C terminal, or both (depending on where the loopy region was located. This was done with the inpaint flavor published in the original RFdiffusion work^14^. 2,000 trajectories were performed each run, followed by the same MPNN and AF2 predictions as above.

## Computational filtering

Precise metrics cutoffs changed for each design campaign to get to an orderable set, but largely focused on interface pAE <10, pLDDT >90, number of hydrogen bonds >11, RMSD < 0.5, sap score <45 and Rosetta ddG < -40^44^ .

## Gene construction of peptide binders

The designed protein sequences were optimized for expression in *E. coli*. Linear DNA fragments (eBlocks, Integrated DNA Technologies) encoding design sequences included overhangs suitable for Golden Gate cloning into LM670 vector (Addgene #191552) for protein expression in *E.coli*. LM670 is a modified expression vector containing a Kanamycin resistance gene, a ccdB lethal gene between BsaI cut sites, and a C-terminal hexahistidine, commonly referred to as His tag.

## Binding screening by Bio-layer interferometry (BLI) or co-lysis of binder and target peptide

For screening for all designs except the ones of partial diffusion design for Amylin-68n(Fig.2a), the designs were screened by BLI (method details described in below relative description). Linear gene fragments encoding binder design sequences were cloned into LM670 using Golden Gate assembly.Golden Gate subcloning reactions of peptide binders were constructed in 96-well PCR plates in 4µL volume. 1µL reaction mixtures were then transformed into a chemically competent expression strain (BL21 (DE3)). After 1 hour recovery in 100 µL SOC medium, the transformed cell suspensions were directly transferred into a 96-deep well plate containing 900 µL of LB media with Kanamycin. After overnight incubation in 37 °C, 100 μL of growth culture were inoculated into 96-deep well plates containing 900 µL of auto-induction media (autoclaved TBII media supplemented with Kanamycin, 2mM MgSO4, 1X 5052). After overnight incubation (6 hours at 37 °C followed by additional 18 hours at 30°C), cells were harvested by centrifugation (15 min at 4000 x g). Bacteria were lysed for 15 minutes in 200 μL lysis buffer (1x BugBuster (Millipore#70921-4), 0.01 mg/mL DNAse, 1 tablet of pierce protease inhibitor tablet/50 mL culture). Lysates were clarified by centrifugation at 4000 g for 10 minutes, before purification on Ni-charged MagBeads (genscript #L00295; wash buffer: 25 mM Tris pH 8.0, 300 mM NaCl, 30 mM Imidazole; elution buffer: 25 mM Tris pH 8.0, 300 mM NaCl, 400 mM Imidazole). Subsequently, the elutions were directly subjected to a BLI test and the final concentration is approximately 1 μM. The designs exhibiting binding signals were subsequently analyzed by BLI through titration.

For Amylin-68n, the designs from partial diffusion were expressed and purified using the same way as mentioned above. In addition to the designs, plasmids expressing target peptide fused with sfGFP (no His tag) were transformed into BL21 (DE3) cells, and overnight outgrowths were cultured in 5 mL of LB media with Kanamycin. After overnight incubation in 37 °C and 250 rpm, growth cultures were inoculated into 50 mL auto-induction media. After overnight incubation in 37 °C and 250 rpm, cells were harvested by centrifugation (15 min at 4000 x g), then resuspended in 20 mL lysis buffer (25 mM Tris-HCl, 150 mM NaCl, 0.1 mg/mL lysozyme, 10 μg/mL DNAse I, 1 mM PMSF). 100 µL of lysate of each binder were mixed with 100 µL of lysate of target peptide fused with sfGFP and incubated at room temperature for 15 min for co-lysis and target binding to the binders. Mixed lysates were applied directly to a 100 µL bed of Ni-NTA agarose resin in a 96- well fritted plate equilibrated with a Tris wash buffer. After sample application and flow through, the resin was thoroughly washed, and samples were eluted in 200 µL of a Tris elution buffer containing 300 mM imidazole. All eluates were sterile filtered with a 96-well 0.22µm filter plate (Agilent 203940-100) prior to size exclusion chromatography. Protein binders were then analyzed for target binding via sfGFP co-elution with the His-tagged binder. High-performance liquid chromatography (HPLC) analyses were conducted using an Agilent HPLC system (<PRODUCT name<). Co-lysates were run on a Superdex200 Increase 5/150 GL column (Cytiva 28990945) with buffer of 25 mM Tris-HCl, 150 mM NaCl. To assess the binding interaction between the target and the binder, we monitored the elution profile of sfGFP using an absorbance wavelength of 395 nm, alongside a simultaneous measurement at 280 nm for total protein content to determine the extent of overlap between 395 nm and 280 nm, which indicates the binding interaction.

## Medium scale protein expression and purification *E.coli* for hits from screening

For further validation, the initial hits were expressed at 50 mL scale via autoinduction for approximately 24 hours, in which the first 6 hours cultures were grown at 37 °C and the remaining time at 22 °C. Cultures were harvested at 4000 g for 10 minutes and resuspended in approximately 20 mL lysis buffer (25 mM Tris-HCl, 150 mM NaCl, 0.1 mg/mL lysozyme, 0.01 mg/mL DNAse, 1 mM PMSF,1 tablet of pierce protease inhibitor tablet/50 mL culture). Sonication was performed with a 4-prong head for 5 minutes total, 10 s pulse on-off at 80% amplitude. The resulting lysate was clarified by centrifugation at 14000 g for 30 minutes. Lysate supernatants were applied directly to a 1 mL bed of Ni-NTA agarose resin equilibrated. After sample application and flow through, the resin was thoroughly washed, and samples were eluted by an elution buffer containing 400 mM imidazole. After elution, protein samples were filtered and injected into an autosampler-equipped Akta pure system on a Superdex S75 Increase 10/300 GL column at room temperature. The SEC running buffer was 25mM Tris-HCl, 150mM NaCl pH 8. Protein concentrations were determined by absorbance at 280 nm using a NanoDrop spectrophotometer (Thermo Scientific) using their extinction coefficients and molecular weights obtained from their amino acid sequences.

## Bio-layer interferometry (BLI) binding experiments

BLI experiments were performed on an Octet Red96 (ForteBio) instrument, with streptavidin coated tips (Sartorius Item no. 18-5019). Buffer comprised 1X HBS-EP+ buffer (Cytiva BR100669) supplemented with 0.1% w/v bovine serum albumin. Prior to target loading, each design was tested for binding against unloaded tips. 50 nM of biotinylated target protein was loaded on the tips for 50 s followed by a 60 s baseline measurement. After loading, all designs underwent a 60 s baseline, 300 s association and 200 s dissociation. Baseline measurements of unloaded tips were subtracted from their matched measurement of the loaded tip. The hits were taken forward for further titration experiments where concentration, association and dissociation times were chosen based on apparent affinity from the single point screen. Global kinetic fitting was used to determine KDs across the dilution series.

## Circular dichroism (CD) experiments

For CD experiments, designs were diluted to 0.4mg/ml in 25 mM Tris-HCl and 150 mM NaCl. Spectra were acquired on a JASCO J-1500 CD Spectrophotometer. Thermal melt analyses were performed between 25 °C and 95 °C, measuring CD at 222 nm. All reported measurements were acquired within the linear range of the instrument.

## Affinity enrichment of Amylin analyzed by LC-MS/MS

### Bead preparation

Anti-amylin binder-coated beads were prepared by conjugating each amylin-targeted binder (Amylin-68n) to paramagnetic M280 Tosylactivated beads (Invitrogen, MA, USA). Each sample reaction conjugated 1 µg of binder to 225 µg of beads. Beads were blocked with a solution of 0.01% bovine serum albumin (BSA) in 0.2 M Tris to minimize non-specific interactions. An off-target binder-conjugated bead was included for quantification of non-specific binding. A BSA-blocked bead without a bound binder was used as a negative control and an anti-GPVGPSGPPGK (GPVG) peptide monoclonal antibody-conjugated bead was used as a positive control for the affinity binding step.

### Sample preparation

Human amylin peptide (non-amidated) was purchased from Anaspec (Fremont, CA, USA) and reconstituted to 2 mg/mL in dimethylsulfoxide (DMSO). A secondary peptide stock (diluted into 50 µM in 5% acetonitrile, 0.1% formic acid, 0.01% BSA in water) was reduced with dithiothreitol (10 mM final concentration) and alkylated with iodoacetamide (30 mM final concentration). Excess iodoacetamide was quenched with additional dithiothreitol (5 mM final added concentration). This solution was diluted to a working stock of 10 μM with dilution solvent. Aliquots of the working stock were made in 1.5 mL LoBind tubes and stored at -20°C to avoid repeated freeze/thaw cycles.

### Human specimens

Human plasma samples were composed of pooled de-identified leftover clinical samples obtained from the clinical laboratories at the University of Washington Medical Center. The use of de-identified leftover clinical samples was reviewed by the University of Washington Human Subjects Division (STUDY00013706).

### Affinity enrichment

Amylin capture experiments were performed using three types of coupled beads (Amlin-68n, off-target binder, BSA-blocked) in phosphate-buffered saline (PBS) containing 0.1% 3-((3- cholamidopropyl) dimethylammonio)-1-propanesulfonate (CHAPS) as well as pooled normal human EDTA-anticoagulated plasma.

Samples were prepared by spiking the working stock of alkylated amylin to a final concentration of 20 nM in 100 µL of either PBS-CHAPS or pooled plasma. Additional PBS-CHAPS was added to each sample, followed by coupled beads. GPVG peptide and anti-GPVG monoclonal antibody-conjugated beads were added to each sample as a positive control. The mixtures were shaken for 1 hr at 900 rpm and room temperature (Thermomixer, Eppendorf, Framingham MA). The supernatant was removed and the beads were washed twice with 200 μL of PBS-CHAPS. Bound peptides were eluted in 50 µL of elution solvent (20% acetic acid, 10% acetonitrile, 10% DMSO, 0.001% BSA in water) with shaking for 8 min (900 rpm, room temperature). Each bead type (two anti-amylin binders, one off-target binder, one BSA-blocked) was assessed in separate samples and each was prepared in triplicate.

Sample analysis was performed by liquid chromatography-tandem mass spectrometry using a Shimadzu Nexera LC-XR HPLC (Columbia, MD, USA) coupled to a Sciex 6500+ triple quadrupole tandem mass spectrometer (Framingham, MA, USA) in multiple reaction monitoring (MRM) mode. Specifications for the liquid chromatography, mass spectrometer, and MRM methods are included in Supplementary Tables x, x, and x.

## Data analysis

Data processing was performed with Skyline Daily (version 23.1.1.459). Chromatographic peak area was calculated by summing the peak area of all transitions for each peptide. The chromatographic peak areas observed during blank (elution solvent) injections were subtracted as background from sample peak areas before performing further data reduction. Signal from BSA and GPVG beads were for quality control of the assay and evaluated prior to processing of the experimental data.

Seven types of samples were analyzed:

1. Group A: Alkylated amylin peptide spiked directly into elution solvent served as the reference peak area for 100% recovery of amylin peptide.
2. Group B: Paramagnetic tosyl-activated beads conjugated to an off-target binder were incubated in PBS-CHAPS spiked with alkylated amylin. The peak area of this negative control was used to quantify nonspecific binding.
3. Group C: Amylin-targeted binders conjugated to paramagnetic tosyl-activated beads were incubated in PBS-CHAPS spiked with alkylated amylin. The peak areas of these samples were used to quantify the percent recovery of amylin by affinity enrichment.
4. Group D: An off-target binder conjugated to paramagnetic tosyl-activated beads was incubated with unspiked plasma. The peak area of this negati/ve control was used to quantify the nonspecific signal from beads binding to plasma components.
5. Group E: Amylin-targeted binders conjugated to paramagnetic tosyl-activated beads were incubated with unspiked plasma. The peak areas observed in these samples were used to quantify the nonspecific signal from the binders binding to plasma components (i.e., assuming no non-amidated amylin in normal plasma).
6. Group F: An off-target binder conjugated to paramagnetic tosyl-activated beads was incubated with spiked plasma. The peak area of this negative control was used to quantify nonspecific binding.
7. Group G: Amylin-targeted binders conjugated to paramagnetic tosyl-activated beads were incubated with spiked plasma. The peak areas of these samples were used to quantify percent recovery of amylin by affinity enrichment.

The percent recovery of each binder-coated bead type was calculated using the following equations:

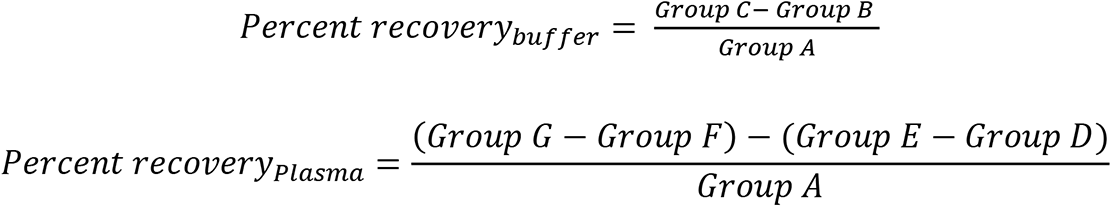

### Preparation of SSM libraries

We performed SSM studies for some of the designed peptide–protein binding pairs to gain a better understanding of the peptide-binding modes, and to search for improved peptide binders. For CP35, we ordered a SSM library covering all the 159 amino acids. The chip synthesized DNA oligos for the SSM library were then amplified and transformed to EBY100 yeast together with a linearized pETCON3 vector. Each SSM library was subjected to an expression sort first, in which the low-quality sequences due to chip synthesizing defects or recombination errors were filtered out. The collected yeast population, which successfully expresses the designed mutants, will be regrown, and subjected to the next round of peptide-binding sorts. Two rounds of with-avidity sorts were applied at 1 μM concentration of C-peptides followed by 1 rounds of without-avidity sorts with C-peptide concentrations at 200nM, 40 nM, 8nM, 1.6nM and 0.32nM. The peptide-bound yeast populations were collected and sequenced using the Illumina NextSeq kit. The mutants were identified and compared to the mutants in the expression libraries. Enrichment analysis was used to identify beneficial mutants and provide information for interpreting the peptide-binding modes. For each mutant, the fraction of cells collected in each of 5 titration sorts of decreasing concentration is measured. The SortingConcentration50 (SC50), the concentration where 50% of the expressing cells are collected, is calculated and plotted in heat maps for the SSM analysis.

## X-ray crystallography

Crystallization experiments were conducted using the sitting drop vapor diffusion method. Initial crystallization trials were set up in 200 nL drops using the 96-well plate format at 20 °C.

Crystallization plates were set up using a Mosquito LCP from SPT Labtech, then imaged using UVEX microscopes and UVEX PS-256 from JAN Scientific. Diffraction quality crystals formed in 0.1M succinic acid, sodium phosphate monobasic monohydrate, glycine mixture at pH 6 and 30% w/v PEG 1000 for Amylin-22. For g3bp1-11 diffraction quality crystals appeared in 0.05 M Calcium chloride dihydrate, 0.1 M BIS-TRIS pH 6.5, and 30% v/v Polyethylene glycol monomethyl ether 550.

Diffraction data was collected at the National Synchrotron Light Source II on beamline 17-ID-1 (AMF) for Amylin-22αβL and Advanced Light Source beamline 821 for g3bp1-11. X-ray intensities and data reduction were evaluated and integrated using XDS^45^ and merged/scaled using Pointless/Aimless in the CCP4 program suite^46^. Structure determination and refinement starting phases were obtained by molecular replacement using Phaser^47^ using the designed model for the structures. Following molecular replacement, the models were improved using phenix.autobuild; with rebuild-in-place to false, and using simulated annealing. Structures were refined in Phenix^48^. Model building was performed using COOT^49^. The final model was evaluated using MolProbity^50^. Data collection and refinement statistics are recorded in Table 5. Data deposition, atomic coordinates, and structure factors reported in this paper have been deposited in the Protein Data Bank (PDB), http://www.rcsb.org/ with accession code 9CC5 and 9CC6.

The highest-resolution shell is shown in parentheses.

### Cell culture

HeLa cells were cultured in DMEM (Gibco, 11965-092) at 37 °C in a humidified atmosphere containing 5% CO2, supplemented with 10% (v/v) FetalClone II serum (Cytiva, SH3006603) and 1% penicillin–streptomycin (ThermoFisher, 15140122).

## Generation of IL2RG-knockout HeLa cells by CRISPR–Cas9 gene targeting

Pooled IL2RG-knockout HeLa cells was generated using the Gene Knockout kit V2 from Synthego, using multi-guide sgRNA targeting IL2RG (guide 1: CAUACCAAUAAUGCAGAGUG, guide 2: UCGAGUACAUGAAUUGCACU and guide 3: GAAACACUGAGGGAGUCAGU). The ribonucleoprotein complex with a ratio of 4.5:1 of sgRNA and Cas9 was delivered following the protocol of the SE Cell Line 4D-Nucleofector^TM^ X Kit S (Lonza, V4XC-1032), using the nucleofection program CN-114 on the Lonza 4D X unit.

## Transient transfection

Plasmids for IL2RG-30-mScarlet, IL2RG-EGFP were synthesized and cloned by Genscript USA, Inc. HeLa cells were seeded at 70–80% confluency in a chambered coverslip with 18 wells (ibidi, 81816). At the same time, HeLa cells were reverse-transfected using Lipofectamine 3000 transfection reagent (ThermoFisher, L3000008) according to the manufacturer’s protocol.

## Fluorescence imaging using 3D structured illumination microscopy

4-color, 3D images were acquired with a commercial OMX-SR system (GE Healthcare). Toptica diode lasers with excitation at 488 nm, and 568 nm were used. Emission was collected on three separate PCO.edge sCMOS cameras using an Olympus 60× 1.42NA PlanApochromat oil immersion lens. 512×512 images (pixel size 6.5 μm) were captured with no binning. Acquisition was controlled with AcquireSR Acquisition control software. Z-stacks were collected with a step size of 250 nm. Images were deconvolved in SoftWoRx 7.0.0 (GE Healthcare) using the ratio method and 200 nm noise filtering. Images from different color channels were registered in SoftWoRx using parameters generated from a gold grid registration slide (GE Healthcare).

## Thioflavin-T (ThT) fluorescence assay

Amylin fibrils at various growth stages (0 h, 3 h and 24 h) were adequately mixed with ThT at molar ratio 1:1 and added into 96-well-plates containing different types and concentrations of binders (Am75, Am36, Am22, Am68n). The samples were then incubated at 37 °C for 1-18 hours with 600 rpm orbital shaking. ThT fluorescence signals were measured using a Thermo Varioskan Flash Multi Detection Microplate Reader (0 h and 3 h) or a Perkin elmer EnSight Multifunctional Microplate Reader (24 h) with excitation wavelength at 440 nm and an emission wavelength at 482 nm.

## Negative-stain electron microscopy (NS-EM) experiment

Samples for negative-stain electron microscopy were dropped onto freshly glow-discharged carbon-coated copper grids and incubated for 1 minute, and excess sample was removed by blotting on filter paper. The grids were then stained with 2 % (w/v) uranyl acetate for 1 minute, and excess uranyl acetate was blotted off. Finally, the grids were examined using a Tecnai Spirit transmission electron microscope (FEI) at an acceleration voltage of 120 kV.

## Acknowledgements

This research used resources (FMX/AMX) of the National Synchrotron Light Source II, a U.S. Department of Energy (DOE) Office of Science User Facility operated for the DOE Office of Science by Brookhaven National Laboratory under Contract No. DE-SC0012704. The Center for BioMolecular Structure (CBMS) is primarily supported by the National Institutes of Health, National Institute of General Medical Sciences (NIGMS) through a Center Core P30 Grant (P30GM133893), and by the DOE Office of Biological and Environmental Research (KP1607011). Amylin fibril inhibition and dissociation work is supported by the Chinese Academy of Sciences (CAS) (XDB37010100), and the basic Research Program Based on Major Scientific Infrastructures, CAS(JZHKYPT-2021-05). We appreciate the help provided by David Juergens in the training of the RFdiffusion model with Joseph L. Watson.

## Author contributions

D.B. directed the work. C.L., K.W. and D.B. designed the research. C.L., H.C., K.W., and H.H. designed, screened and experimentally characterized the binders. J.L.W. developed the sequence input RFdiffusion and secondary structure specification algorithm used for IDP/IDR binder design.

C.L. prepared samples for crystallography, A.K.B., A.K. and E.B. obtained all the crystal structures shown in this manuscript. H.C., C.L., and K.W. performed all-by-all specificity BLI.

W.Y. constructed the SSM library. C.L. screened SSM library and analyzed the SSM result with the help from B.C., D.R.H. and X.W.. H.C. and J.D. experimentally validated the colocalization of IL2RG binders to the target in mammalian cells. S.S. and A.N.H carried out the LC–MS/MS peptide detection. X.Z. and P.Z. performed the Amylin fibrils formation inhibition and Amylin fibrils dissociation experiments. S.R.G., A.M. and M.L. carried out additional scaled-up protein purification. M.B. helped with prion binder design. C.L., K.W., H.C. and D.B. wrote the manuscript with input from the other authors.

## Code availability

Code explanation and examples of binder design using RFdiffusion can be found at *<GITHUB_LINK=.* Partial-diffusion code explanation and examples can be found at *<GITHUB_LINK=*.

## Data Availability

Crystal structures of Amylin-22αβL and G3bp1-11 have been deposited in Protein Data Bank, with the accession IDs 9CC5 and 9CC6, respectively.

